# BATF3-dependent induction of IL-27 in B cells bridges the innate and adaptive stages of the antibody response

**DOI:** 10.1101/2020.06.26.117010

**Authors:** Hui Yan, Rui Wang, Jingwei Wang, Shuai Wu, Maria Fernandez, Carlos E. Rivera, Christian Cervantes, Justin B. Moroney, Xiao-Dong Li, Nu Zhang, Hong Zan, Xiangzhi Meng, Fushun Zhang, Siyuan Zheng, Yidong Chen, Zhinan Yin, Ross Kedl, Booki Min, Christopher A. Hunter, Yan Xiang, Paolo Casali, Zhenming Xu

## Abstract

B cells are exposed to innate and T cell stimuli during the antibody response, although whether and how they functionally integrate such signals are unclear. Here we have identified IL-27 as the cytokine specifically produced by murine B cells upon sequential stimulation by TLR ligands and then CD154 and IL-21, the hallmark factors of T follicular helper cells, and during the T-dependent antibody response to a conjugated hapten or virus infection. B-cell *Il27p28* transcription is concomitant with increased locus accessibility and depends on newly induced BATF3 transcription factor. IL-27-producing B cells are inefficient in antibody secretion, but cooperate with IFNγ to promote proliferation, survival, class-switching and plasma cell differentiation of CD40-activated B cells, leading to optimal IgG2a and IgG1 responses. Overall, IL-27-producing B cells function as “helper” B cells that integrate the innate and adaptive stages of the antibody response.

**One-sentence summary:** B cells integrate innate TLR and adaptive CD40 signals to induce BATF3 transcription factor for production of IL-27, which together with INFg optimizes antibody responses.

## Main Text

For effective antibody responses to infectious and environmental antigens, specific B cells recognize arrayed epitopes that crosslink their B cell receptors (BCRs) and later need to be engaged by CD154 (CD40 ligand, CD40L) expressed on cognate T helper (Th) cells for robust proliferation and differentiation in germinal centers (GCs), from which plasma cells emerge to secrete mature antibodies. These are of switched Ig isotypes (IgG, IgA and IgE) and high affinity, as underpinned by Ig locus class switch DNA recombination (CSR) and somatic hypermutation (SHM), respectively (*1*). Prior to the T-dependent adaptive phase, both specific and bystander B cells have already been exposed to antigen-or host cell-derived stimuli that engage innate receptors, particularly Toll-like receptors (TLRs). Accordingly, engagement of B-cell TLR can synergize with BCR crosslinking to induce AID, a DNA deaminase essential for CSR and SHM (*2*), and elicit class-switched and high-affinity IgG antibodies during the T-independent phase of antibody responses or stand-alone T-independent responses (*3*). In immune-competent subjects, TLR ligands can function as adjuvants to potentiate antibody responses, partially by priming dendritic cells (DCs) that in turn activate Th cells (*4*). However, the relevance of TLR priming to B cell activation by T cell stimuli in the T-dependent antibody response remains unclear.

Independent from their differentiation into antibody-secreting plasma cells, B cells are known to produce cytokines (*5–8*), including those that direct CSR from IgM to specific Ig isotypes, such as IL-4 (IgG1), TGFβ (IgA) and IFNg (murine IgG2a), thereby supplementing cytokine production by Th cells. In addition, mouse B cells can be induced to produce cytokines IL-10 and IL-35 (heterodimer of *Il12a*- encoded IL12α and *Ebi3-encoded* EBI3), thereby acting as immunosuppressive B regulatory (Breg) cells in several pathophysiological conditions (*9, 10*). Here, we explore new regulatory functions of B cells, starting with profiling cytokine gene induction in B cells by innate stimuli TLR ligands in combination with T cell stimuli, thereby mimicking B cell activation during the antibody response.

## Results

### Potent IL-27 induction in B cells

Purified mouse naive spleen B cells expressed high levels of *Il10* when stimulated by TLR4 ligand LPS (**fig. S1A–S1C**). Addition of membrane-bound recombinant CD154 to LPS dampened *Il10* expression and specifically induced transcription of *Il27p28*, but not *Il12a* (**Fig. 1A, 1B, fig. S1C, S1D, Table S1, S2**), leading to intracellular IL-27p28 protein expression and secretion of cytokine IL-27, which is composed of IL-27p28 and EBI3 (**Fig. 1C**, **fig. S1E**) – *Ebi3*, an ortholog of *EBI3* identified in EBV-infected human B cells (*11*), was similarly upregulated. *Il27p28* was also induced when other TLR9 ligands were used and in lymph node B cells, which are mature follicular B cells (**fig. S1F, S1G**).

**Fig. 1.**
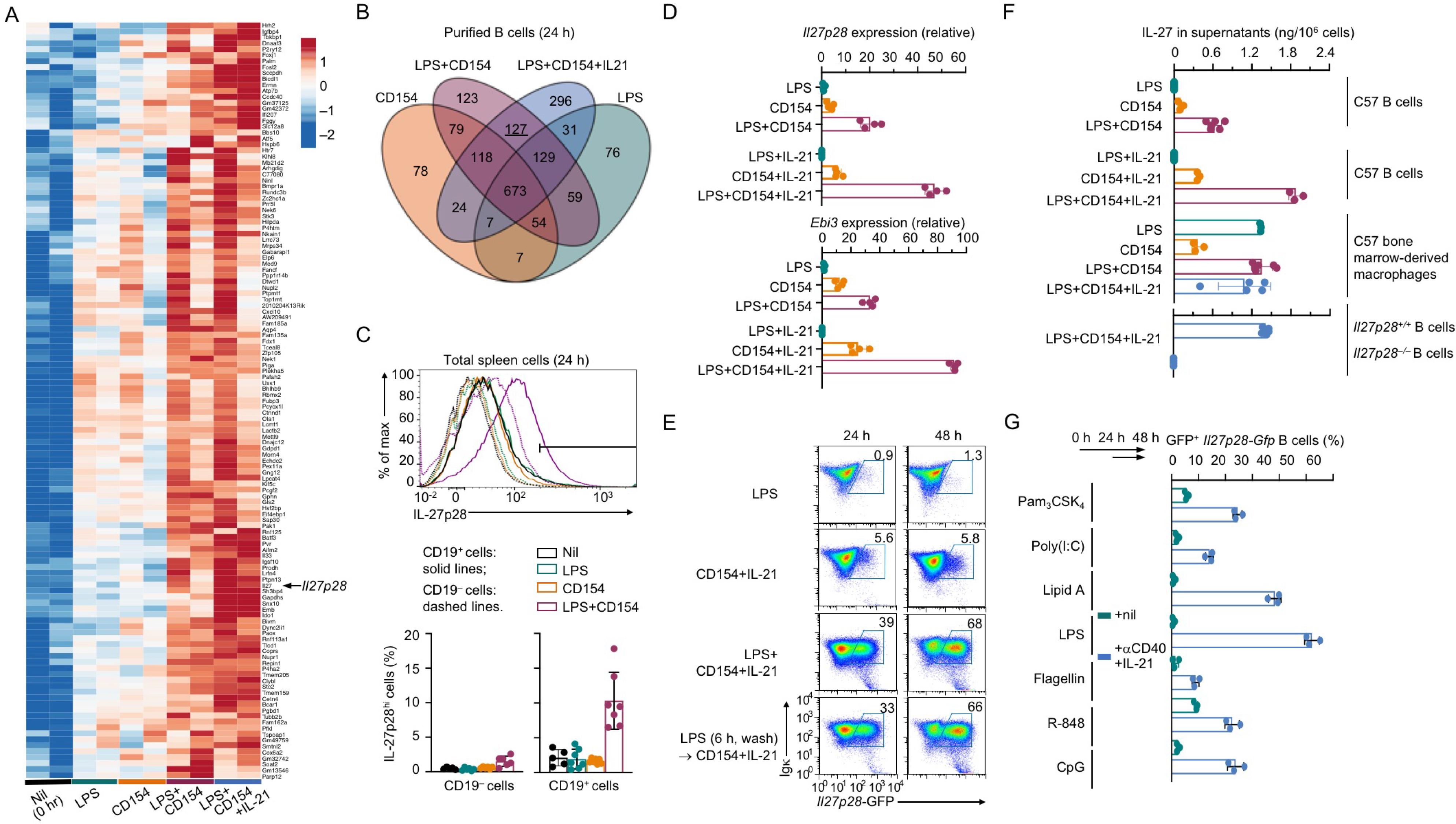
Induction of IL-27 in B cells. **(A,B)** Heatmap and Venn diagram of 127 differentially expressed genes (DEGs) in B cells after stimulation with LPS plus CD154 or LPS plus CD154 and IL-21, but not LPS or CD154, for 24 h. Data are from two independent stimulation and RNA-Seq experiments. Genes expressed in at least one sample were analyzed. **(C)** Intracellular staining of IL-27p28 in CD19^*–*^ and CD19^+^ cells after total spleen cells were stimulated for 48 h. **(D)** qRT-PCR analysis of *Il27p28* and *Ebi3* expression in purified B cells stimulated, as indicated. Data were normalized to *Cd79b* expression and expressed as the ratio to values of freshly isolated B cells (0 h). **(E)** Flow cytometry analysis of GFP expression in purified *Tg(Il27p28-Gfp*) B cells co-stimulated or sequentially stimulated, as the readout of transcription activity in the *Il27p28* locus. Representative of three independent experiments. **(F)** ELISA of IL-27 secreted into the supernatants of B cells after stimulation for 48 h. **(G)** Percentages of GFP^+^ cells by flow cytometry after purified *Tg(Il27p28-Gfp*) B cells were primed with a TLR ligand, as indicated, for 24 h and then stimulated with αCD40 and IL-21 for 24 h.

While marginally affected by IL-4, a Th2 cytokine that synergizes with CD154 in activating B cells, B-cell IL-27 induction was markedly enhanced by IL-21, the hallmark cytokine of T follicular helper (Tfh) cells (**Fig. 1A, 1D, fig. S2A, Table S1**). This resulted in *Il27p28* promoter activation (GFP expression) in 70% of B cells from *Tg(Il27p28-Gfp*) reporter mice and IL-27 secretion at high levels, e.g., comparable to those from LPS-activated macrophages, in which IL-27 production was not enhanced by CD154 or IL-21 (**Fig. 1E, 1F, fig. S2B, S2C**). Priming B cells with LPS, which upregulated surface CD40 expression, followed by stimulation with CD154 and IL-21 was as effective as co-stimulation in inducing IL-27 (**Fig. 1E, fig. S3A, S3B**), in association with enhanced B cell survival (but not expression of selected surface markers), as compared to CD154 and IL-21-stimulated B cells (**fig. S3B–S3D**). Such induction by sequential innate and adaptive stimuli could be extended to priming by other TLR ligands (irrespective of their potency in inducing B cell proliferation, plasma cell differentiation and expression of CD11b and CD11c) and CD40 engagement by an agonistic mAb (αCD40), concomitant with robust cluster formation (**Fig. 1G, fig. S4A–S4C**). BCR crosslinking, by contrast, failed to prime B cells for IL-27 induction (**fig. S4D**), showing the specific synergy of TLR and CD40 signals in this process.

### B cells are important IL-27 producers in vivo

Owing to their ability to induce IL-27p28 and large numbers in secondary lymphoid organs, CD19^+^ B cells upregulated IL-27p28 levels in C57 mice immunized with alum-mixed NP-CGG together with LPS and accounted for 75% of all IL-27p28 expression (**Fig. 2A, fig. S5A, S5B**) – likewise, they contributed to the majority of homeostatic IL-27p28 levels, as also confirmed by highly correlating *Il27p28* and *Ebi3* expression with that of *Cd79b* (**fig. S5C**). Spleen cells displaying IL-27p28 expression were mostly B220^+^ and localized in B cell follicles and extrafollicular areas, with a subset of such cells showing concentrated signals at the plasma membrane (**Fig. 2B, fig. S6A**). IL-27p28^+^ B cells could also be elicited in draining lymph nodes and the spleen upon intranasal (i.n.) infection by vaccinia virus (the Western Reserve strain, VV_WR_), whose DNA activates TLR8, but not TLR9 in DCs (*12*), starting at d 3 and markedly increasing by d 7, and segregated within *Il27p28-GFP^+^* cells in *Tg(Il27p28-Gfp*) reporter mice (**Fig. 2C, fig. S7A, S7B**) – EBI3 expression was also upregulated in *Il27p28-GFP^+^* B cells. Whether induced by NP-CGG plus alum and LPS immunization or VVWR infection, IL-27p28^+^ B cells were distinct from IL-10^+^ B cells, e.g., GFP^+^ B cells in *Il10^IRES-Gfp^* mice (**Fig. 2B, fig. S6B, S7C, S7D**).

**Fig. 2.**
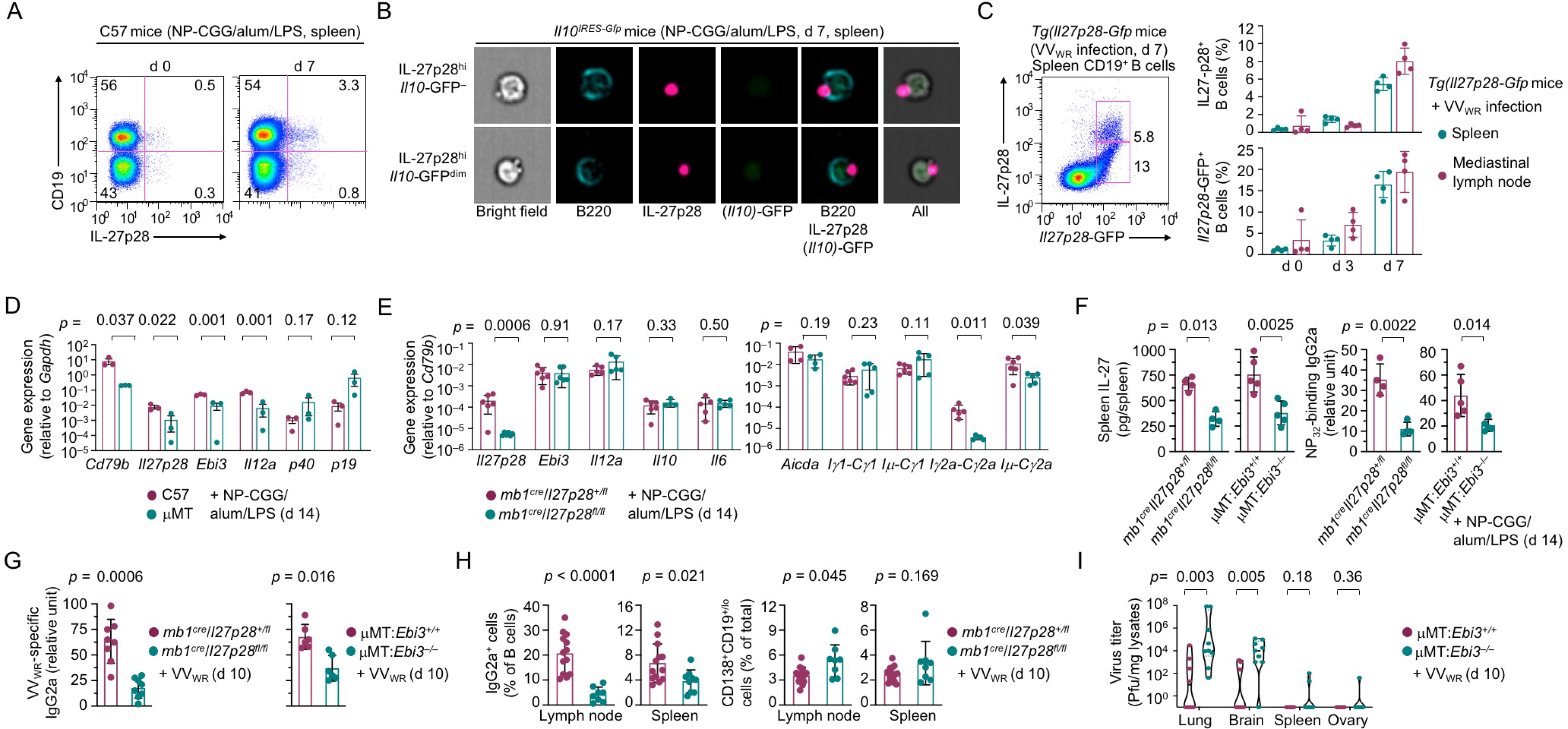
Induction and function of IL-27-producing B cells *in vivo.* **(A)** Intracellular staining of IL-27p28 in CD19^+^ cells in C57 mice before and after immunization, as indicated. Representative of three independent experiments. **(B)** Flow imaging analysis of B220, IL-27p28 and GFP (indicating *Il10* transcription) in immunized *Il10^IRES-Gfp^* mice (representative of three mice in two independent experiments). **(C)** Flow cytometry analysis of IL-27p28 protein and GFP expression to quantify IL-27p28 expression and *Il27p28* gene transcription in the spleen and mediastinal lymph node B cells from *Tg(Il27p28-Gfp*) mice after virus infection. **(D)** qRT-PCR analysis of transcript levels of cytokine genes, as indicated, in the spleen of C57 and μMT mice 7 d after immunization. Data were normalized to *Gapdh* expression. **(E)** qRT-PCR analysis of cytokine gene expression and CSR-related transcripts in the spleen of *mb1^cre^Il27p28^fl/fl^* and *mb1^cre^Il27p28^+/fl^* mice after immunization. Data were normalized to *Cd79b* expression and expressed as the fold of values in *mb1^cre^Il27p28^+/fl^* mice (mean and s.d.). **(F)** ELISA of IL-27 production in the spleen (left panel) and NP-binding IgG2a in the circulation (right panel) in *mb1^cre^Il27p28^fl/fl^*, μMT:*Ebi3^−/−^* and their respective “wildtype” mouse counterparts, as indicated, after immunization. **(G)** ELISA of virus-specific IgG2a in the circulation in *mb1^cre^Il27p28^fl/fl^*, μMT:*Il27p28^−/−^*, μMT:*Ebi3^−/−^* and their respective “wildtype” mouse counterparts 10 d after VV_WR_ infection. **(H)** Quantification of IgG2a^+^ B cells and CD138^hi^ plasma cells/plasmablasts in mediastinal lymph nodes and the spleen by flow cytometry in VV_WR_-infected *mb1^cre^Il27p28^fl/fl^* and *mb1^cre^Il27p28^+/fl^* mice. **(I)** Quantification of VV_WR_ titers in different organs in VV_WR_-infected μMT:*Ebi3*^+/+^ and μMT:*Ebi3*^−/−^ mice.

In μMT mice, which had virtually no B cells, expression of *Il27p28* and *Ebi3* was significantly decreased, leading to reduced IL-27 secretion – expression of *Il10* and *Il12a* was also impaired (**Fig. 2D, fig. S8A–S8C**). In NP-CGG plus alum and LPS immunized *mb1^+/cre^Il27p28^fl/fl^* mice (**fig. S9A**), mixed bone marrow chimera mice generated using μMT and *Il27-p28^−/−^* mice as donors (μMT:*Il27p28^−/−^*, **fig. S9A, S9B**) or chimera mice using μMT and *Ebi3^−/−^* mice as donors (μMT:*Ebi3*^−/−^, **fig. S9C**), ablating B-cell *Il27p28* or *Ebi3* resulted in B cells-specific loss of IL-27p28 or EBI3 expression, reduction in their gene expression in the spleen and defective IL-27 secretion into the circulation – IL-27 levels peaked at d 14 in “wildtype” mice (**Fig. 2E**, **2F, fig. S9D, S9E**). The higher IL-27p28 and EBI3 levels in B cells that those in CD19^-^ cells also confirmed an important role of B cells in IL-27 production (**fig. S9D**).

### B-cell IL-27 optimizes IgG2a responses

In association with their defective IL-27 production, mice with B cell-specific deficiency in *Il27p28 (mb1^+/cre^Il27p28^fl/fl^*) or *Ebi3* (μMT: *Ebi3^−/−^*) were impaired in mounting IgG2a responses to immunization with NP-CGG plus alum and LPS, showing reduced levels of NP-binding IgG2a, including high-affinity NP_4_-binding IgG2a at d 7 and NP_30_-binding IgG2a at d 14, but largely normal levels of NP-binding IgG1, IgG2b and IgG3 (**Fig. 2F**, **fig. S10A**). Total IgG2a as well as IgM and IgG3, but not IgG1 or IgG2b, were also reduced (**fig. S10B**), likely reflecting a role of B-cell IL-27 in Ig responses to the CGG and LPS components in the immunogen. The defects in IgG2a antibody responses were due to impairment in CSR to IgG2a, as shown by significant reduction in germline *Iγ2a-Cγ2a* and post-recombination *Iμ-Cγ2a* transcripts, the molecular marker of ongoing and completed CSR to IgG2a, respectively, and class-switched IgG2a^+^ B cells (**Fig. 2E, fig. S11A**). By contrast, B cell survival and differentiation into GC B cells and plasma cells were not affected in these mice; neither was expression of AID (**Fig. 2E, fig. S11B**).

Murine IgG2a and human IgG3 are the IgG isotype elicited first upon viral infections (*13, 14*). IgG2a has potent anti-viral activities through complement activation, engagement of activating Fcγ receptors and binding of viral antigens via both F(ab) arms due to their long-hinge region for effective neutralization – by contrast, IgG1 appears later and effects neutralization by high-affinity antigen-binding via a single F(ab). Accordingly, *mb1^+/cre^Il27p28^fl/fl^* and μMT:*Ebi3^−/−^* mice produced less virus-specific IgG2a (and IgG) antibodies upon i.n. VV_WR_ infection (**Fig. 2G, fig. S10C**), consistent with their reduced CSR to IgG2a (**Fig. 2H**). This resulted in persistence of viruses in μMT:*Ebi3^−/−^* mice in the lung, dissemination of such viruses into the brain, and loss of more body weight (**Fig. 2J, fig. S11D**). Thus, IL-27-producing B cells play an important role in CSR to IgG2a and optimal IgG2a responses, including those that effect the host anti-viral immunity.

### Requirement for BATF3 in IL-27^+^ B cell generation

Concomitant with IL-27p28 induction in B cells, the chromatin accessibility in the promoter and a proximal enhancer (Enhancer I) of the *Il27p28* locus was increased, as compared to LPS-stimulated B cells, in which two upstream enhancers (Enhancer II and III) were also accessible (**Fig. 3A, fig. S12A, Table S3**). To identify *trans*-factors recruited to such differentially accessible regions (DARs), transcriptome analysis of B cells stimulated to express *Il27p28* at different levels revealed that *Il27p28* expression showed a strong positive correlation with that of *Batf3*, the best among all transcription factor genes and top 5 among all genes (R^2^=0.799; **Fig. 3B, Table S4, Table S5**) – *vice versa*, expression of *Batf3* tightly correlated with that of *Il27p28* among all cytokine and chemokine-coding genes (**fig. S12B, Table S6, Table S7**). Importantly, DARs in IL-27^+^ B cells were enriched with the motif bound by the BATF family of factors (**Fig. 3C, Table S8**), among which BATF3 was induced in such B cells, in contrast to the constitutive *Batf* expression and no *Batf2* expression in B cells (**fig. S12C, Table S5**).

**Fig. 3.**
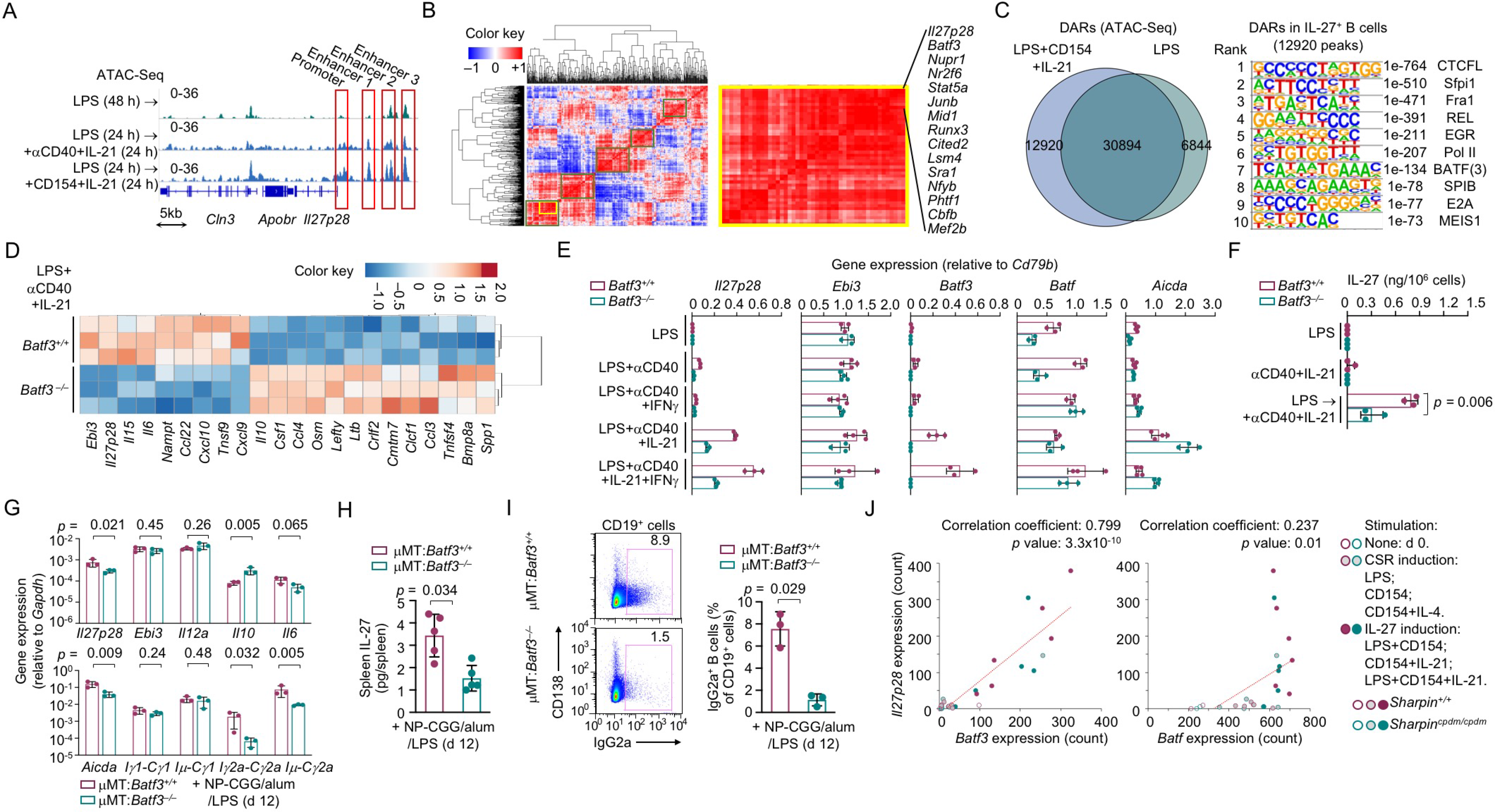
BATF3 mediates in IL-27 induction in B cells. **(A)** ATAC-Seq analysis of the chromatin accessibility of the *Il27p28* locus in stimulated B cells. **(B)** RNA-Seq and correlation analysis of the expression level of *Il27p28* and that of 441 expressed transcription factors in duplicate B cells stimulated *in vitro* with nil (0 h), (CSR-inducing) LPS, CD154 and CD154 plus IL-4, as well as (IL-27-inducing) LPS plus CD154, CD154 and IL-21, and LPS plus CD154 and IL-4, for 24 h. (**C**) Venn diagram depiction of DARs in B cells stimulated with LPS plus CD154 and IL-21, as compared to those in LPS-stimulated B cells (left) and motifs enriched in such DARs (right). (**D**) Heatmap of cytokine-encoding genes differentially expressed in stimulated *Batf3^+/+^* and *Batf3^−/−^* B cells. (**E,F**) qRT-PCR analysis of gene expression in *Batf3^+/+^* and *Batf3^−/−^* B cells stimulated for 48 h (**E**) and ELISA of IL-27 secretion by these cells (**F**). (**G-I**) qRT-PCR analysis of gene expression (**G**), ELISA of IL-27 in the spleen (**H**) and flow cytometry analysis of switched IgG2a^+^ B cells (**I**) in μMT:*Baf3^+/+^* and μMT:*Batf3^−/−^* mice immunized with NP-CGG plus alum with LPS for 12 d. (**K**) Correlation analysis of *Il27p28* expression with that of *Batf3* or *Batf* in *Sharpin^+/+^* and *Sharpin^cpdm/cpdm^* B cells stimulated to induce CSR or IL-27 production, as indicated (*Batf2* was not expressed in B cells in any condition).

*Batf3^−/−^* B cells were severely impaired in *Il27p28* transcription induction and IL-27 secretion, exemplifying an overall defect in expressing genes involved in the IFN*γ* and/or anti-viral responses (**Fig. 3D–3F, fig. S13A, S13B, Table S9-S11**). They, on the other hand, showed heightened expression of *Il10* and genes involved in myeloid cell activation/differentiation, as characteristically upregulated by LPS (**fig. S13B, Table S12**), suggesting the cytokine producing profile in TLR-primed B cells was dictated by BATF3 (if induced) without affecting the overall chromatin accessibility (**fig. S13C**). Mice with B cell-specific *Batf3* deficiency (μMT:*Batf3^−/−^*) showed decreased *Il27p28* expression and IL-27 secretion, but increased *Il10* expression, upon immunization with NP-CGG plus alum and LPS (**Fig. 3G, 3H**). They were impaired in mounting specific IgG2a responses, concomitant with reduced generation of IgG2a^+^ B cells and expression of Iγ2a-Cγ2a and Iμ-Cγ2a transcripts as well as *Aicda* (**Fig. 3G–3I, fig. S14**), showing a non-redundant B cell-intrinsic role of BATF3 in IL-27 induction and IgG2a responses.

### Supporting role of NF-κB in IL-27 production

BATFs belong to the leucine zipper transcription factor family, which also includes the AP-1 factor c-JUN and c-FOS. With a high-affinity DNA-binding domain but no transactivation domain, they modulate gene expression by competing with c-JUN or c-FOS for the same AP-1 sites to inhibit transcription or by interacting with (positive or negative) transcription factors recruited to neighboring *cis-*elements to strengthen the activity of such partner factors (*15–17*). In addition to BATF-binding sites, κB sites were enriched in DARs in IL-27-producing B cells (**Fig. 3C, Table S8**). Accordingly, the *Il27p28* promoter region and three enhancers have consensus κB motifs near an AP-1 site, with three putative BATF/κB composite sites in Enhancer I and III (**fig. S15**). In B cells stimulated to induce BATF3 and IL-27p28, the promoter region was bound by the NF-κB p65 subunit, which was synergistically activated (as indicated by Ser536 phosphorylation, p65-S536p) by LPS and αCD40 (**fig. S16A, S16B**). In such B cells (as well as in LPS-stimulated B cells), genome-wide binding sites of NF-κB p65 were focused on promoters and proximal enhancers, as clustering around transcription start sites (TSSs; **fig. S16C**). They were also enriched in BATF(3)-binding motifs (**fig. S16D**), suggesting corecruitment of NF-κB and BATF3 to the same *cis-*elements.

Engagement of CD40, which is a TNFR superfamily member, elicited stronger B-cell NF-κB p65 activation than TLR signals (**Fig. S15B**). Consistent with the dependency of B-cell CD40 (and TNFR in general) on **l**inear **ub**iquitin **a**ssembly **c**omplex (LUBAC, composed of HOIL, HOIP and SHARPIN) for signaling (*18, 19*), p65-S536p was reduced in *Sharpin^cdmp^* B cells, in which a single nucleotide deletion causing frameshift and premature translation termination, resulting in no SHARPIN expression (**fig. S17A**). These B cells were defective in *Il27p28* and *Ebi3* expressing and IL-27 secretion (**fig. S17B**). Irrespective of their *Sharpin* genotype, B cells had highly concordant expression of *Il27p28* and *Batf3*, the two most upregulated genes by IL-27-inducing stimuli (**Fig. 3K, fig. S17C, S17D**).

### Synergy of B-cell IL-27 and IFNγ receptor signals in vivo

As the target of B cell-produced IL-27, IL-27Ra, which is the IL-27-specific subunit of the receptor, was highly expressed in B cells – the other subunit GP130 is shared by receptors of IL-6 family cytokines (**fig. S18A, S18B**). In *Cd19^+/cre^ Il27ra^fl/fl^* mice and mixed bone marrow chimera μMT:*Il27ra^−/−^* mice, specific IgG2a response to immunization with NP-CGG plus alum and LPS was reduced (**Fig. 4A, fig. S18C–S18E, fig. S19A, S19B**), in agreement with previous studies (*20*). Accordingly, CSR to IgG2a was impaired in μMT:*Il27ra^−/−^* mice (**fig. S19C–S19G**). Upon VV_WR_ infection, μMT:*Il27ra^−/−^* mice produced less virus-specific IgG2a and failed to eradicate virus in the lung or limit virus spreading, consistent with their reduced serum virus-neutralization activities (**Fig. 4B–4F**).

**Fig. 4.**
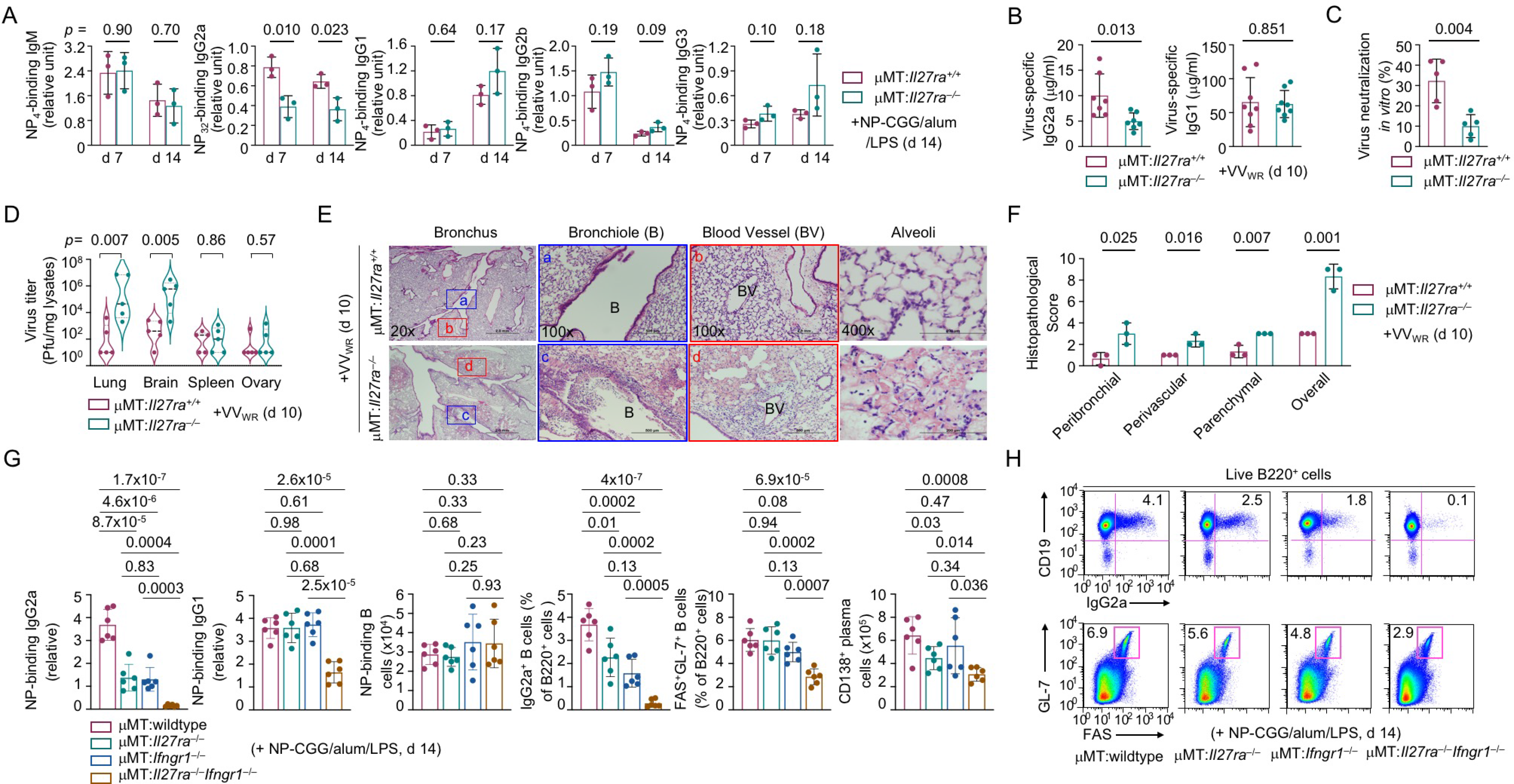
Function of IL-27 receptor signals in B cells. **(A)** ELISA of NP-binding IgM and IgG isotypes in μMT:*Il27rα^−/−^* and μMT:*Il27rα^+/+^* mice immunized with NP-CGG plus alum with LPS. **(B-F)** ELISA of virus specific IgG2a and IgG1 in serum samples in VV_WR_-infected μMT:*Il27rα^−/−^* and μMT:*Il27rα^+/+^* mice (**B**), the virus-neutralizing activity of serum samples (**C**), virus titers in organs (**D**), and lung histopathology (**E**) and scores (**F**). **(G,H)** ELISA of NP-binding IgG isotypes and FACS analysis of B cells in mixed bone marrow chimera mice with B cell-specific double and single KO of *Il27ra* and *Ifngr1*, as indicated, after immunization with NP-CGG, alum and LPS.

While significantly reduced (by 60%) upon ablation of IL-27^+^ B cells or *Il27ra*-expressing B cells as their potential target cells, IgG2a responses, as well as CSR to IgG2a, were abrogated in mice with B cell-specific compound deficiency in *Il27ra* and *Ifngr1* (μMT:*Il27ra^−/−^ Ifngr1^−/−^*; **Fig. 4G, 4H, fig. S20A**). The complementary roles of B-cell IL-27 and IFN*γ* receptor signals would provide a parsimonious explanation for IgG2a elimination in whole-body knockout (KO) *Il27ra^−/−^Ifngr1^−/−^* mice (**fig. S20B**). They, together with possible reciprocal induction of IL-27 and IFNγ in all cell sources, would also explain the 90% reduction in IgG2a levels in whole-body single KO *Il27ra^−/−^* and *IfngrΓ~* mice (**fig. S20B**), as compared to the 60% reduction in B cell-specific single KO μMT:*Il27ra^−/−^* and μMT:*Ifngr1^−/−^* mice.

Unexpectedly, μMT:*Il27ra^−/−^Ifngr1^−/−^* mice were defective in the IgG1 response to immunization, concomitant with impairment in the generation of GL-7^+^FAS^+^ GC B cells and CD138^+^ plasma cells as well as Tfh cells (**Fig. 4G, 4H, fig. S20A**). Accordingly, whole-body KO *Il27ra^−/−^Ifngr1^−/−^* mice displayed reduced homeostatic IgG1 levels, in addition to the abolished IgG2a levels, and early mortality upon VV_WR_ infection (**fig. S20B, S20C**). *Ebi3^−/−^Ifng^−/−^* mice had similar defects (**fig. S20D**), showing the synergy of IL-27 and IFN*γ* in effecting anti-viral responses by B cells and CD8^+^ T cells (*21, 22*).

### IL-27 and IFNγ receptor signals modulate B cell differentiation

Consistent with the synergy their respective receptor signals in B cells *in vivo*, IL-27 and IFNγ synergized to activate STAT1, but did not activate any other STATs analyzed, in CD154-primed B cells (**fig. S21A**). These cells could also be conditioned by IL-21 to display more STAT1 activation in response to IL-27 or IFNγ, which together, however, abrogated activation of STAT3 by IL-21 (**fig. S21A**). Such divergent activation of different STATs was associated with differential regulation of many gene sets, as exemplified by IL-27 and IFNγ upregulation of “Type 1” genes (particularly *Tbx21, Cxcr3, Cxcl9* and *Cxcl10*) and germline Iγ2a-Cγ2a transcription while downregulating germline Iγ1-Cγ1 transcription and *Aicda* expression (**Fig. 5A, 5B, fig. S21B, Table S13, S14**). IL-27 enhanced plasma cell differentiation in B cells stimulated with CD154 plus IL-21 and, together with IFNγ, rescued such B cells from the killing effect of IL-21 (**Fig. 5C, fig. S21C, S21D**). Nevertheless, IL-27 and IFNγ failed to direct CSR to IgG2a in B cells stimulated with CD154 or CD154 plus IL-21, consistent with lack of robust *Aicda* expression in either case (**Fig. 5D, fig. S21B**).

**Fig. 5.**
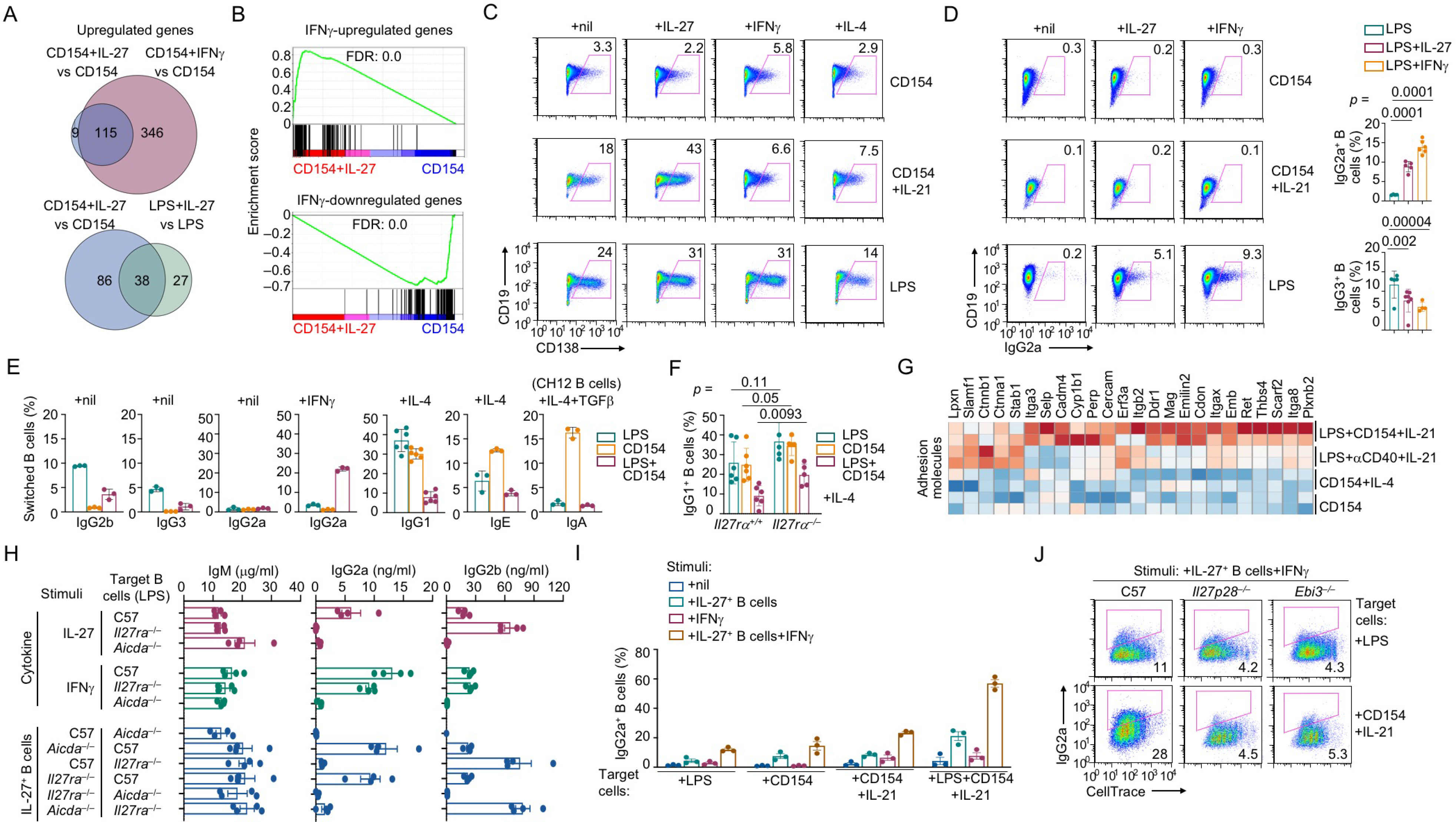
Synergy of IL-27^+^ B cells and IFNγ on target B cell differentiation. **(A)** Venn diagram depiction of DEGs induced by IL-27 and IFNγ in B cells stimulated with CD154 for 24 h (top) or DEGs induced by IL-27 in B cells stimulated with CD154 or LPS for 24 h (bottom). **(B)** GSEA analysis of DEGs upregulated (top) or downregulated (bottom) by IL-27 and IFNγ in B cells stimulated with CD154 for 24 h. **(C,D)** Flow cytometry analysis of plasma cell differentiation (**C**) and CSR to IgG2a (**D**) in B cells stimulated, as indicated, for 96 h. **(E)** Flow cytometry analysis of CSR to different Ig isotypes in B cells stimulated, as indicated, for 96 h. **(F)** Flow cytometry analysis of CSR to IgG1 in *Il27ra^+/+^* and *Il27ra^−/−^* B cells stimulated, as indicated, for 96 h. **(G)** Heatmap of adhesion molecule-encoding genes upregulated in IL-27^+^ B cells (top two rows), as compared to B cells stimulated with CD154 plus nil or IL-4 (bottom rows). **(H)** ELISA of IgM, IgG2a and IgG3 secreted by *Aicda^−/−^*, C57 and *Il27ra^−/−^*B cells, as target B cells, stimulated with LPS for 96 h in the presence of cytokine IL-27 or IFNγ (top 2 clusters), or IL-27^+^ B cells (bottom cluster), as generated from B cells of different genotypes (as indicated) and then co-cultured with target B cells for 72 h. **(I)** FACS analysis of CD45.2^+^ target B cells labeled with CellTrace™ (to track proliferation) and activated, as indicated, 16 h before co-cultured for 72 h with nil, CD45.1^+^ IL-27^+^ B cells, IFNγ or both CD45.1^+^ IL-27^+^ B cells and IFNγ, as indicated (n=3). **(J)** FACS analysis in CD45.1^+^ target B cells, as indicated, 16 h before co-cultured with CD45.2^+^ B cells of different genotypes that had been stimulated with LPS plus αCD40 and IL-21 for 36 h as inducing B cells plus IFNγ. Representative of two independent experiments.

In contrast to T-dependent “primary” stimulus CD154 (*1*), alone or with IL-21, the T-independent primary stimulus LPS programmed B cells to undergo CSR to IgG2a and concomitant reduction of CSR to IgG3 in response to IL-27 or IFNγ despite effect of IFNγ in killing TLR-activated B cells, with little impact on plasma cell differentiation (**Fig. 5C, 5D, fig. S21C, S21E**). These effects were associated with IL-27 and IFNγ activation of STAT1 and their additive effect in STAT3 activation – with IL-27 and IFNγ also synergized with IL-4 to induce moderate levels of STAT5 activation (**fig. S21A**). They were likely underpinned by the induction of *Aicda* and germline Iγ2a-Cγ2a transcription together with *Tbx21* and other Type 1 genes (**Fig. 5A, fig. S21B, Table S15**). Finally, IL-27, but not IL-35, antagonized the effect of IL-4 in LPS-stimulated B cells by downregulating induction of Type 2 gene expression and CSR to IgG1 without affecting B cell proliferation or survival (**fig. S22, Table S16, S17**).

### IL-27^+^ B cells synergize with IFNγ to induce CSR to IgG2a

Owing to their production of IL-27 that inhibited CSR to non-IgG2a isotypes, LPS and CD154-stimulated B cells underwent little CSR to IgG2b, IgG3 or – in the presence of IL-4 or TGFβ – IgG1, IgE or IgA (**Fig. 5E, 5F, fig. S23A–S23C**). IL-27^+^ B cells were also refractory to plasma cell differentiation and IgM secretion, consistent with reduced expression of *Prdm1*, which encodes BLIMP1, the master transcription factor of plasma cell differentiation (**fig. S23D–S23F**). *Batf3* and *Sharpin*, while important for IL-27 induction, were dispensable for CSR and plasma cell differentiation (**fig. S24**).

B cells stimulated to produce IL-27 failed to switch to IgG2a, but could to do so in the presence of IFNγ (**Fig. 5E, fig. S25A, S25B**). They upregulated selected adhesion molecules and formed doublets with LPS-stimulated B cells (**Fig. 5G, fig. S25C, Table S18**). They also induced IgG2a switching and secretion in target B cells activated by LPS, and remained IgG2a^-ve^ *per se* (**Fig. 5H, fig. S25D**). In target B cells activated by CD154 or αCD40 (alone or with IL-21), IL-27^+^ B cells – unlike recombinant IL-27 – induced CSR to IgG2a (**Fig. 5I, fig. S26A**). By contrast, “control” B cells stimulated with LPS was inefficient in inducing IgG2a in target B cells (**fig. S26B**). In addition, IL-27^+^ B cells and IFNγ together enhanced survival and proliferation of target B cells activated by CD154 and IL-21 (**fig. S26C**).

The ability of IL-27^+^ B cells to direct T-dependent CSR to IgG2a was synergistically enhanced by IFNγ (but not IL-4), leading to switching in 25% and 50% of target B cells activated by CD154 and αCD40, respectively, plus IL-21 in a manner dependent on IL-27p28 and EBI3 (**Fig. 5I, 5J, fig. S26A**). Such a synergy was independent of the boost of target B cells proliferation, as the increase was across cell divisions, with early divisions also displaying high IgG2a levels (**fig. S26B**). Finally, the synergistic IgG2a induction was maximized (over 60%) when target B cells were activated by LPS plus CD154 and IL-21, the same stimuli that also induced IL-27^+^ B cells (**Fig. 5I, fig. S26**), showing that IL-27^+^ “regulatory” B cells could be induced to differentiate into IgG2a-expressing “effector” B cells *in trans*.

## Discussion

Here, we have reported the identification of IL-27-producing B cells, their induction and function in the T-dependent antibody response, and underlying mechanisms. These include the pivot role of BATF3 in controlling the B cell cytokine production program and a new paradigm of induction of CSR to IgG2a, the IgG isotype immensely relevant to the host defense against viral infections.

Since its discovery (*23*), IL-27 has been thought to be produced mainly by monocytes. As unveiled here, IL-27^+^ B cells could be robustly induced by a unique condition, i.e., priming by TLR signals (which can likely be substituted by other innate signals) and then exposure to adaptive CD40 signals, and potentiated proliferation, survival and differentiation of target B cells in response to T cell stimuli, thereby bridging the innate phase and adaptive phase of the T-dependent IgG2a and IgG1 responses. TLR ligands, whether co-encapsulated with antigens in nanoparticles or co-delivered with them as separate entities (*4, 24*), enhance IgG2a and IgG1 responses by acting directly on B cells (and DCs). They would allow B cells to induce AID/CSR upon BCR engagement to effect T-independent antibody responses or readily produce IL-27 upon engagement by Tfh cells to boost T-dependent responses. Absent such adaptive signals from BCR or Tfh cells, TLR-primed B cells may secrete IL-10, IL-6 or even IL-35, thereby inhibiting the antibody response, as previously suggested (*10*). In TLR-licensed B cells, BATF3, upon induction by Tfh cell factors, skewed the B cell cytokine program in favor of IL-27 and away from IL-10. Such opposing effects could be underpinned by the activities of BATF3 as both a transcription activator and inhibitor, as previously shown also in CD4^+^ T cells (*25*). The marginal impact of BATF3 on CSR highlights its difference and BATF, which – like EBI3 – was originally identified in EBV-transformed human B cells (*26*) and promotes CSR (*27*), but is partially redundant at best with BATF3 in IL-27 induction. Likewise, the role of SHARPIN in NF-kB activation for IL-27 induction, but not CSR, contrasts the role of RAB7 in promoting NF-kB activation for CSR induction (*28*).

TLR priming would allow both antigen-specific and bystander B cells to receive Tfh cell help, most likely by upregulating CD40 expression. This can tighten the CD40:CD154 engagement and sustain the “entanglement” between B cells and Tfh cells, thereby strengthening the MHC II:TCR interaction involving cognate B cells or making up for it in bystander B cells – upregulated surface CD40 would be internalized upon engagement by CD154 (*28*). IL-27^+^ B cells, despite losing the ability in secreting antibodies, produced IL-27 that may in turn act on Tfh cells to upregulate IL-21 (*29*), consistent with a putative B cell function in governing Tfh cell dynamics in GCs, in addition to providing ICOSL (*30*). This also raises the possibility of a positive feedback loop involving IL-21^+^ Tfh cells and IL-27^+^ B cells to nearly synchronize IL-21 and IL-27 production for the antibody response to viral infections, after an earlier DC-produced IL-27 peak that would mobilize CD8^+^ T cells for rapid anti-viral responses – IL-27 may also be dependent on BATF3 may also mediate production of IL-27 in DCs, within which it is recruited to Enhancer II and III in the *Il27p28* locus (ChIP-Seq data from Ref. (*31*)).

As helper cells and likely distinct from age-associated B cells or T-bet^+^ B cells (*32*), IL-27^+^ B cells, through secreted IL-27 and likely also other elements, such as adhesion molecules involved in “synapsis”, promoted the growth of target B cells activated by T-dependent stimuli and their differentiation into effector plasma cells that produced large amounts of IgG2a and IgG1 antibodies. Such enhancement effects were reciprocally reinforced, complemented and/or backed up by those of a powerful collaborator in IFNγ, most notably CSR to IgG2a at a high level (up to 50%), which rivals the high efficiency of CSR to IgG1 induction. IFNγ may be derived from Th1 CD4^+^ T cells or type 1 Tfh cells, leading to a cytokine milieu rich in IL-21, IL-27 and IFNγ and essential for CSR to IgG2a – within this milieu, even IL-27^+^ B cells can potentially differentiate into IgG2a^+^ B cells. IL-27^+^ B cells may be engaged in tri-cellular interactions with CD40-activated target B cells and IL-21^+^IFNγ^+^ Tfh cells to strengthen the GC response, consistent with an important role of IFNγ (and STAT1) in antibody and autoantibody responses (*33, 34*).

The importance of such cellular interactions is also emphasized by the inability of recombinant IL-27 and IFNγ in the induction of CSR to IgG2a. Finally, IL-27 and IFNγ collaborate to mediate host responses to viral infections and likely the generation and function of Tregs and ILCs (*35–37*).

The effect of IL-27 (as produced by BATF3^+^ B cells *in vivo*) and IFNγ in optimizing the response of their target B cells to T cell stimuli is likely underpinned by the unique pattern of STAT signal outputs, with IL-21 playing a key role. IL-21 is critical for GC B cell as well as *Prdm1* induction and plasma cell differentiation in a STAT3-dependent manner (*38, 39*), but promotes B cell death, as previously reported and re-emphasized here (*40*). While IL-27 or IFNγ-mediated STAT1 activation was enhanced by IL-21, IL-21-activated STAT3 is synergistically abrogated by IL-27 and IFNγ. The different effects on STAT activation would ultimately differentially affect the target genes of IL-27 and IFNγ. The impact of abrogated STAT3 activation on *Prdm1* expression and plasma cell differentiation may be mitigated by STAT1 and other transcription factors. As AID promotes GC B cell death by generating double-strand DNA breaks (DSBs) that, if overwhelming the DNA repair machinery, can instigate apoptosis (*41, 42*), downregulation of this genotoxic enzyme by IL-27 and IFNγ likely contribute to their pro-survival effect on target B cells, as assisted by the upregulation of the anti-apoptotic genes. That IL-4 and IL-21 together trigger massive death of CD154-stimulated B cells suggests that these two cytokines do not function on the same B cell *in vivo* without strong B-cell survival factors (such as BAFF or combined IL-27 and IFNγ), consistent with their different induction kinetics and the notion that most CSR to IgG1 completes before the GC reaction initiates (*43–45*). Nevertheless, plasma cell differentiation of IgG1^+^ B cells would still dependent on the full-blown GC reaction and be defective in μMT:*Il27a^−/−^ Ifngr1*^−/−^ mice. Overall, IL-27^+^ B cells, together with IFNγ, coordinate innate and adaptive immune elements, either on the cell surface or secreted, to regulate antibody responses and likely diverse pathophysiological conditions owing to pleiotropic functions of IL-27 as both a pro-inflammatory and anti-inflammatory cytokine (*46*).

## Acknowledgments

We wish to thank: Dr. Zhijie Liu for providing anti-NF-kB p65 rabbit polyclonal Ab, Dr. Elizabeth Leadbetter for *Il10^IRES-Gfp^* mice, Dr. William Kaiser for *Sharpin^cpdm/cpdm^* mice, and Mr. Daniel Chupp for technical help (UTHSCSA); and Dr. Joseph Craft for advice (Yale).

## Author contributions

Conceptualization (H.Y., R.W. and Z.X.), investigation (H.Y., R.W., J.W., S.W., M.F., C.E.R., C.C., J.B.M., X.-D. L., N.Z., X.M., F.Z. and Z.X.), resources (R.K., B.M., C.A.H. and Y.X.), visualization (H.Z., S.Z. and Y.C.), funding acquisition (P.C. and Z.X.) and supervision (Z.X.).

## Data and materials availability

All materials are available upon request, pending material transfer agreements. Deep-sequencing data (RNA-Seq, ATAC-Seq and ChIP-Seq) are being deposited to the NCBI. All data, code, and materials are available to any researcher for purposes of reproducing or extending the analysis.

## Supplementary Materials

Materials and Methods

Figures S1–S26

Tables S1-S21

Supplementary References and Notes (*47–49*)

## Materials and Methods

### Mouse strains

C57BL/6J (C57, JAX, stock #000664), *Ebi3^−/−^* (#008691), *Batf3^−/−^* (#013755), *Il27ra^−/−^* (#018078), *Ifng^−/−^* (#002287), *Ifngr1^−/−^* (#003288), *Sharpin^cpdm/cpdm^* (#007599), *Il10^IRES-Gfp^* (#008379), μMT (#002288) and CD45.1^+^ (*Ptprc^a^*, #002014) were originally from the Jackson Laboratory. *Il27p28^fl/fl^, Tg(Il27p28-Gfp*) and *Il27ra^fl/fl^* mice were as described (*47–49*). *Il27p28^fl/fl^* mice were bred with *mb1^+/cre^* mice (but not *Cd19^+/cre^* mice, as the *Il27p28* and *Cd19* genes are <200 Kb apart on the Chromosome 7) to generate B cell-specific KO *mb1^+/cre^Il27p28^fl/fl^* mice, and *Il27ra^fl/fl^* mice with *Cd19^+/cre^* mice to generate *Cd19^+/cre^Il27ra^fl/fl^* mice. C57BL/6N-A^tm1Brd^ *Il27^tm1a(EUCOMM)Wtsi)^* homozygotic mice, as re-derived from frozen sperms (EMMA Mouse Repository, strain # 10189), were confirmed to be unable to express *Il27p28* transcripts and secrete IL-27 by qRT-PCR and ELISA, respectively, due to the interference of gene expression by the large *LacZ-Neo^r^* selection cassette inserted between Exon 1 and 2 (**fig. S9A**). Hence, they were used as *Il27p28^−/−^* mice. They were also bred with a C57 mouse strain that expressed Flipase under the control of the *Actb* locus (JAX, stock #003800) to delete the cassette, thereby yielding a mouse line carrying the *Il27^tm1c^* allele, as equivalent to the *Il27p28^fl^* allele used in this study (**fig. S9A**).

To generate mice with B cell-specific deficiency in *Il27p28, Ebi3, Batf3* or *Il27ra* using the mixed bone marrow chimera mouse approach, age- and sex-matched recipient C57 mice from the same breeding batch in JAX were treated with neomycin sulfate (2 mg/ml in drinking water) for one week before being g-irradiated with a lethal dose (1000 rad, or 10 Gy, 12.5 min) via a cesium source. After 24 h, mice were randomized into two groups and injected through the tail vein 2.5×10^6^ mixed bone marrow cells, as comprised of 80% (2×10^6^) cells from a single μMT mouse and 20% (0.5×10^6^) cells from either a KO mouse or its age- and sex-matched C57 mouse counterpart (“wildtype”). Prior to mixing, bone marrow cells isolated form tibia and fibula of donor mice were depleted of T cells by incubation with biotinylated anti-CD3 mAb and Magnisort^®^ Streptavidin Negative Selection Beads (eBioscience). Chimeric mice were monitored by flow cytometry analysis of CD45, CD19, CD4, CD8, CD11b, CD11c and/or Gr1 expression of mononuclear cells in the circulation for at least 6 weeks to confirm the immune system reconstitution. To verify B cell-specific gene deficiency in mice, CD19^+^CD4^-^CD8^-^, CD19^-^CD4^+^CD8^-^ and CD19^-^CD4^+^CD8^-^ peripheral blood cells were sorted (BD FACSAria™ II) for genomic DNA extraction and genotyping using PCR involving specific primers. All mice were maintained in a pathogen-free vivarium of the University of Texas Health Science Center at San Antonio (UTHSCSA) and were used for experiments at 8-12 weeks of age, without any apparent infection or disease. While female mice were used in most experiments, in the experiments in which male mice were also used, they did not display significant differences from female mice in the same group.

### Mouse immunization and vaccinia virus infection

For immunization, mice were injected intraperitoneally (i.p.) with 100 μg of NP-CGG (in average 16 molecules of NP, 4-hydroxy-3-nitrophenyl acetyl, conjugated to one molecule of CGG, chicken γ-globulin; Biosearch Technologies) in the presence of 100 μl of alum (Imject^®^ Alum adjuvant, ThermoFisher) in the central-left abdomen area and 10 μg of LPS (from *E. coli*, serotype 055:B5 and deproteinized; Sigma-Aldrich) in 100 μl of PBS in the central-right abdomen area. For intranasal route (i.n.) infection of the wildtype Western Reserve (VV_WR_) strain of vaccinia virus (VV), 5×10^4^ plaque forming units (Pfu) of infectious viral particles (in 20 μl PBS) were delivered into the nostrils (10 μl each) in infect mice, as anesthetized by ketamine and xylazine (50 mg/Kg and 5 mg/Kg of body weight, respectively). Infected mice were kept on a warm pad until waking up and being transferred to original cages. Each mouse was weighed one day before infection and daily after infection until moribund (e.g., losing 25% or more of the initial body weight) or sacrificing at predetermined end-points. All protocols were in accordance with the rules and regulations of the Institutional Animal Care and Use Committee (IACUC) of UTHSCSA.

### Vaccinia virus

HeLa cells (ATCC^®^ CCL2.2™) were maintained in monolayer in 150cm^2^ tissue culture flask in DMEM supplemented with fetal bovine serum (FBS, 10% v/v, Invitrogen) and penicillin-streptomycin/amphotericin B (1% v/v). At the time of infection, 10^9^ cells growing at the log phase were infected with VV_WR_ virus according to a standard protocol at an m.o.i. of 1 for 72 h at 37°C in the presence of DMEM and FBS. Infected cells were pelleted and resuspended in 14 ml of buffer (10 mM Tris-Cl, pH 9.0). After homogenization in a glass Dounce homogenizer, nuclei were removed by centrifugation (900 g, 5 m, 5°C). The supernatant was sonicated and layered onto a cushion of 17 ml of 36% sucrose. After centrifugation (32,900g, 80 min, 4°C), the virus pellet was resuspended in 1 ml of 1 mM Tris-Cl (pH 9.0). After titration, viruses were stored in 100 μl aliquots at −80°C until thawed to be used only once for infection.

For titration of virus titers in tissues, a portion of the lung, spleen as well as brain and ovary (two target organs) isolated from infected mice were weighed and homogenized in 1 ml PBS. The homogenates were 10-fold serially diluted and added to Vero cells (ATCC^®^ CCL-81™) grown at the log-phase in 6-well plates in the presence of DMEM supplemented with 1% FBS and 0.5% (w/v) methylcellulose (Sigma-Aldrich). After 2 d of incubation at 37°C, the medium was removed and cells were washed once before addition of crystal violet (0.1%) for incubation at RT for 1 h. After drying, the number of plaques was counted for quantification.

For virus neutralization assay, purified VV_WR_ was mixed with serum samples from virus-infected mice at 4°C for 1 h (in a total volume of 1 ml). For each neutralization condition, three independent mixtures were set up and inoculated onto monolayers of Vero cells in 6-well plate. After incubation at 37°C for 1 h, the inoculum was replaced with 2 ml DMEM supplemented with 1% FBS and 0.5% (w/v) methylcellulose. The plates were incubated at 37°C for 2 d before being processed for plaque counting in the same way as the virus titration analysis.

### Mouse B cell isolation and purification

Mouse leukocytes were isolated from single cell suspensions prepared from the spleen, mediastinal lymph node, and pooled axillary, inguinal and cervical lymph nodes using a 70-μM cell strainer. Lymph node cells were collected in RPMI 1640 medium (Invitrogen) supplemented with FBS (10% v/v, Invitrogen), penicillin-streptomycin/amphotericin B (1% v/v) and 50 μM β-mercaptoethanol (RPMI-FBS) and resuspended in PBS for purification. Spleen cells were resuspended in ACK Lysis Buffer (Lonzo) to lyse red blood cells and, after quenching with RPMI-FBS, were resuspended in PBS for purification. To prepare leukocytes from mouse peripheral blood, freshly collected blood (100 μl) were treated with heparin (5,000 U) to avoid aggregation and then subjected to red blood cell lysis using ACK Lysis Buffer (100 μl) twice and suspended with RPMI-FBS.

To purify B cells, spleen and lymph node cells were subjected to negative selection (against CD43, CD4, CD8, CD11b, CD49b, CD90.2, Gr-1 or Ter-119) using the EasySep™ Mouse B cell Isolation Kit (StemCell™ Technologies). Cell preparations were stained with fluorophore-conjugated antibodies specific for CD45 or CD45.2, CD19, CD4, CD8, CD11b, CD11c and/or Gr1 at 4°C for 30 m and verified by FACS analysis to contain >95% CD19^+^CD11b^-^Gr1^-^CD11c^-^ B cells and no CD4^+^ or CD8^+^ T cells. For further purification, these B cell preparations were sorted (BD FACSAria II™) to yield 99% of CD19^+^CD11b^-^Gr1^-^CD11c^-^ cells. To purify CD19^+^ B cells by positive selection, single cell suspensions containing 5×10^6^ splenocytes were incubated with 2 μg biotinylated anti-CD19 mAb before reacted with Magnisort^®^ Streptavidin Positive Selection Beads (eBiosciences) following manufacturer’s instructions.

### In vitro B cell culture

To culture mouse B cells purified from the spleen or lymph nodes, or total splenocytes, cells were cultured (3×10^5^ cell/ml, 2 ml in general in a 24-well plate) in RPMI-FBS with the following stimuli at the dose indicated unless specified otherwise: CD154 (3 U/ml), agonistic anti-CD40 mAb (clone FGK4.5, 5 μg/ml), LPS (3 μg/ml), TLR1/2 ligand Pam_3_CSK_4_ (100 ng/ml, Invivogen), TLR4 ligand lipid A (1 μg/ml, Invivogen), TLR7 ligand R-848 (30 ng/ml, Invivogen), TLR9 ligand ODN1826 (sequence 5’-TCCATGACGTTCCTGACGTT-3’) with a phosphorothioate backbone (CpG, 1 μM; Eurofins), F(ab’)2 of a goat anti-mouse μ chain antibody (anti-μ F(ab’)2, 1 μg/ml; Southern Biotech), which crosslinks IgM BCR, or anti-Igδ mAb (clone 11-26c conjugated to dextran, anti-δ/dex, 100 ng/ml; Fina Biosolutions), which crosslinks IgD BCR, alone or in combination. CD154 was prepared as membrane fragments isolated from Sf21 insect cells infected by CD154-expressing baculovirus, as we described (*2, 28*) – membrane fragments from non-infected Sf21 cells failed to activate B cells to proliferate. Other stimuli added to the culture included recombinant IL-21 (2 ng/ml), IL-27 (100 ng/ml, R&D Systems), IFNγ (30 ng/ml) and IL-4 (4 ng/ml), at the indicated dose unless specified otherwise. In sequential stimulation experiments, B cells were cultured in the presence of the first stimulus (or stimuli) for 24 h before the second stimulus (or stimuli) were added to the culture.

In B cell co-culture experiments, B cells prepared from CD45.1^+^ or CD45.2^+^ mice were stimulated with LPS, CD154 (or αCD40) and/or IL-21 for 36 h, washed twice with RPMI-FBS to remove the stimuli and resuspended in RPMI-FBS before used as stimulating B cells. Target B cells were labeled with CellTrace™ Yellow and activated for 16-24 h by stimuli, as indicated, before mixed with stimulating B cells (CD45.2^+^ or CD45.1^+^) at the 1:1 ratio (3×10^5^ cells/ml each), alone or with other cytokines, and cocultured for additional 72 h before FACS analyses. Stimulating and target B cells were also co-cultured at different ratios for different time periods, as indicated, before doublet formation analysis.

### Bone marrow derived macrophage differentiation and stimulation

Bone marrow cells (10^7^) isolated from tibia and fibula were cultured for 10 d in RPMI-FBS in the presence of conditioned media from L-929 cells that express macrophage colony-stimulating factor (which promotes differentiation into macrophages), resulting in the generation of CD11b^+^ cells in over 90% of all cells. After stimulation with LPS (3 μg/ml or as indicated), CD154 (3 U/ml or as indicated) and/or IL-21 (2 ng/ml), for 24 h, supernatants were collected for ELISA of IL-27 and cells for RNA extraction and transcript analysis.

### Flow cytometry and intracellular staining

To analyze B cells stimulated *in vitro*, cells were harvested, stained with fluorochrome-conjugated mAbs in Hank’s Buffered Salt Solution plus 0.1% BSA (BSA-HBSS) for 20 m and analyzed for Igg3, Igg1, Igg2b and Igg2a expression (CSR to IgG3, IgG1, IgG2b and IgG2a), CD138 (plasma cell marker) and other B cell surface molecules on a flow cytometer (BD LSR II™ or Celesta™). To analyzed immune cells *in vivo*, spleen or blood cells (2×10^6^) were first stained with fluorophore-labeled mAbs to surface markers in the presence of mAb Clone 2.4G2, which blocks FcgIII and FcgII receptors, and FVD-506. After washing, cells were resuspended in HBSS for FACS.

To analyze expression of IL-27p28 and EBI3, cells (2×10^6^ cells) were incubated with Brefeldin A (100 μM) in RPMI-FBS (2 ml) at 37 °C for 4 h. After pelleting, cells were stained with fluorophore-labeled mAbs to surface markers and FVD. After washing, cells were resuspended in the BD Cytofix/Cytoperm™ buffer (250 μl) and incubated at 4°C for 20 m. After washing twice with the BD Perm/Wash™ buffer, cells were counted again and 10^6^ cells were resuspended in 100 μl of BD Cytofix/Cytoperm™ buffer for staining with APC- or PE-Cy7-conjugated anti-IL27p28 mAb, PE-Cy7-conjugated anti-EBI3 mAb or isotype-matched control IgG Abs at 4°C for 30 m. After washing with BD Perm/Wash™ buffer, cells were analyzed by FACS. All data were analyzed by FlowJo^®^ (BD).

### B cell proliferation and survival

For flow cytometry analysis of B cell proliferation *in vivo*, mice were injected twice i.p. with 2 mg of bromodeoxyuridine (BrdU) in 200 μl PBS, with the first and second injection at 24 h and 20 h prior to sacrificing, respectively. Spleen B cells were analyzed for BrdU incorporation (by anti-BrdU mAb staining) and DNA contents (by 7–AAD staining after cells were permeabilized) using the APC BrdU Flow kit^®^ (BD) following the manufacturer’s instructions. For B cell proliferation analysis *in vitro*, B cells were labeled with CellTrace™ CFSE or CellTrace™ Yellow (ThermoFisher) following the manufacturer’s instructions and then cultured for 96 h. Cells were analyzed by flow cytometry for CellTrace™ CFSE or CellTrace™ Yellow intensity, which was reduced by half after completion of each cell division, as CellTrace™ CFSE or CellTrace™ Yellow-labeled cell constituents were equally distributed between daughter cells. The number of B cell divisions was determined by the FlowJo^®^ software to calculate the average cell divisions that B cells had completed. To analyze survival of *in vitro* stimulated B cells, cells were collected and stained with anti-CD19 Ab and FVD or 7-AAD to distinguish live cells (FVD^-^ or 7-AAD^-^) from cells undergoing apoptosis/necrosis.

### ELISA

To determine titers of total IgM, IgG1, IgG2a, IgG2b and IgG3, sera were first diluted 4-to 128-fold with PBS (pH 7.4) plus 0.05% (v/v) Tween-20 (PBST), and cell culture supernatants were diluted 2-to 10-fold with PBS. Two-fold serially diluted samples and standards for each Ig isotypes were incubated in 96-well plates pre-treated with sodium carbonate/bicarbonate buffer (pH 9.6) and coated with pre-adsorbed goat anti–IgM or anti–IgG (to capture IgG1, IgG2a, IgG2b and IgG3) Abs (all 1 mg/ml). After washing with PBST, captured Igs were detected with biotinylated anti–IgM, –IgG1, – IgG2a, –IgG2b or –IgG3 Abs, followed by reaction with horseradish peroxidase (HRP)-labeled streptavidin (Sigma-Aldrich), development with o-phenylenediamine (OPD) and measurement of absorbance at 492 nm. Ig concentrations were determined using Prism^®^ (GraphPad) or Excel.

To analyze titers of antigen-specific antibodies, sera were diluted 100-to 1,000-fold in PBST. Twofold serially diluted samples were incubated in a 96-well plate pre-blocked with BSA and coated with NP32-BSA (in average 32 NP molecules on one BSA molecule, to capture all NP-specific antibodies) or NP_4_-BSA (in average 4 NP molecules on one BSA molecule, to capture high-affinity NP-specific antibodies). Captured Igs were detected with biotinylated Ab to IgM, IgG1, IgG2a, IgG2b or IgG3. Data are relative values based on end-point dilution factors or area under curve (AUC).

ELISA of IL-27 in the spleen or circulation *in vivo* or secreted into the supernatants of cultured B cells *in vitro*, total spleen lysates, serum samples or supernatants were analyzed by the Mouse IL-27 ELISA Read-Set-Go^®^ Kit (Invitrogen) following the manufacturer’s’ instructions. Briefly, samples were diluted two-fold with the diluent and incubated in 96-well plates pre-coated with an anti-IL-27p28 polyclonal Ab. After washing, samples were reacted with a biotin-conjugated anti-EBI3 polyclonal Ab followed by the color development using TMB substrate. IL-27 was quantified using a standard curve derived from the reading of serially diluted purified recombinant IL-27, with a sensitivity of 16 pg/ml.

### ELISPOT

For ELISPOT analysis of total IgM^+^ and IgG1^+^ antibody secreting cells (ASCs) as well as NP-binding IgM^+^ and IgG1^+^ ASCs, Multi-Screen^®^ filter plates (Millipore) were activated with 35% ethanol, washed with PBS and coated with pre-adsorbed goat anti–IgM or anti–IgG, or NP4–BSA (5 μg/ml) in PBS. Single spleen or bone marrow cell suspensions were cultured at 50,000 cells/ml in RPMI-FBS at 37°C for 16 h. After supernatants were removed, plates were incubated with biotinylated goat anti-mouse IgM or anti-IgG1 Ab, as indicated, for 2 h and, after washing, incubated with HRP-conjugated streptavidin. Plates were developed using the Vectastain AEC peroxidase substrate kit (Vector Laboratories). The stained area in each well was quantified using the CTL Immunospot software (Cellular Technology) and depicted as the percentage of total area of each well for ASC quantification.

### Flow imaging and immunofluorescence microscopy

Splenocytes from immunized *Il10^IRES-Gfp^* mice were first stained for surface B220 and then intracellular IL-27p28 before resuspended in BSA-HBSS at the concentration of 2×10^7^ cells/ml. They were analyzed using an ImageStream X multispectral imaging flow cytometer (Amnis^®^), with >10,000 events collected in each sample. Images were analyzed using IDEAS image-analysis software (Amnis^®^), with single-stained samples being used for compensation of fluorescence signals. To visualize IL-27p28 expression and GCs, spleens were embedded in OCT (Tissue-tek) and snap-frozen on dry ice. Cryostat sections (7 μm) were fixed in pre-chilled acetone at −80°C for 30 m and air dried at 25°C. Sections were stained with fluorochrome-conjugated Abs at 25°C for 1 h in a moist chamber. After washing with HBSS, sections were mounted using ProLong^®^ Gold Anti-Fade Reagent (Invitrogen) and examined under a Zeiss LM710 confocal microscope.

### RNA isolation, qRT-PCR and RNA-Seq

Purified B cells (2×10^6^) cultured and stimulated *in vitro* or freshly isolated and purified by positive selection (see above) were pelleted and total RNA was extracted using the RNeasy Mini Kit (Qiagen). For *ex vivo* analysis of gene expression in the spleen, 1/6 of spleen freshly isolated from each mouse was immediately preserved in the RNAlater™ stabilization reagent until cDNA synthesis. First-strand cDNA was synthesized from 1 μg of RNAs suing the Superscript™ III system (Invitrogen) and qPCR was performed using SYBR Green (Dynamo HS kit; New England Biolabs) kit in a QuantStudio™ 3 Reak-Time PCR Instrument (Applied Biosystems). The ΔΔCt method was used to analyze levels of transcripts and data were normalized to the levels of *Gapdh* or as indicated.

For RNA-Seq, after RNA integrity was verified using an Agilent Bioanalyzer 2100 (Agilent), RNA was processed using an Illumina TruSeq RNA sample prep kit v2 (Illumina). Clusters were generated using TruSeq Single-Read Cluster Gen. Kit v3-cBot-HS on an Illumina cBot Cluster Generation Station. After quality control procedures, individual RNA-Seq libraries were pooled based on their respective 6-bp index portion of the TruSeq adapters and sequenced at 50 bp/sequence using an Illumina HiSeq 3000 sequencer. Resulting reads, typically 20 million reads per sample, were checked by assurance (QA) pipeline and initial genome alignment (Alignment). After sequencing, demultiplexing with CASAVA was employed to generate a Fastq file for each sample. All sequencing reads were aligned with the (GRCm38/mm10) reference genome using HISAT2 default settings, yielding Bam files, which were then processed using HTSeq-count to obtain counts for each gene. RNA expression levels were determined using GENCODE annotation. Differential expression analysis was performed using the Deseq2 package in R post-normalization based on a Benjamini-Hochberg false discovery rate (FDR)-corrected threshold for statistical significance of *p_adj_* <0.05 or raw *p* value <0.01. Transcript read counts were transformed to ln(x+1) used to generate heatmaps in Clustvis. Volcano plots depicting log_2_-fold change and raw or adjusted *p* values were generated in R.

### Assay for Transposase-Accessible Chromatin using Sequencing (ATAC-Seq)

For ATAC-Seq library preparation, Omni-ATAC, which minimizes mitochondrial DNA contamination, was used to increase the library complexity and the signal-to-noise ratio. Briefly, stimulated B cells were centrifuged at 500 g for 5 min at 4°C, washed with 50 μl cold PBS and centrifuged again at 500 g for 5 min at 4°C. The cell pellet was then resuspended in 50 μl cold lysis buffer (10 mM Tris pH 7.4, 10 mM NaCl, 3 mM MgCl_2_, 0.1% NP-40, 0.1% Tween-20, 0.01% digitonin), incubated on ice for 3 min, quenched with 500 μl cold Tween-only-buffer (10 mM Tris pH 7.4, 10 mM NaCl, 3 mM MgCl_2_, 0.1% Tween-20). After pelleting by centrifugation (500g, 30 min at 4°C), nuclei were immersed in tagmentation reaction mixture (5 μl 5X insertion buffer, 2 μl Tn5-50 transposase, 8 μl sterile PBS, 10 μl nuclease-free H_2_O) at 37°C for 45 min, resulting in a 30 μl mixture. After dilution in 30 μl of 2X nuclei lysis/deactivation buffer (300 mM NaCl, 100 mM EDTA, 0.6% SDS, 1.6 μg Proteinase K) and incubation for 30 min at 40°C, DNA was purified using Qiaquick columns (Qiagen) and amplified using Illumina-compatible index primers to prepare the libraries, which were normalized through qPCR analysis. DNA was enriched for fragments <1000 bp using Zymo Select-a-Size DNA Clean and Concentrator MagBead kit according to manufacturer’s instructions. After the quality check on an Agilent Bioanalyzer 2100, samples were pooled at an equimolar ratio for 50-bp single-end run on a HiSeq 3000. For sequence analysis, ATAC-Seq data sets were aligned to the mouse genome (GRCm38 /mm10) by Bowtie2 with the default parameters. Uniquely mapped reads were masked using Samtools. Peaks were identified by MACS2 with FDR (q-value 0.05) comparing specific ATAC reads with unspecific control reads derived from sequencing matching input samples without building the shifting model and set extension size 200 bp. Peaks were annotated by annotatePeaks.pl (-d option) in HOMER using the peak file. The unique peaks of each sample were identified by MAnorm 1.3.0. For motif enrichment analysis, unique peaks was selected for extraction of 150 bp centered on the summit, and subjected to motif analysis using HOMER.

### Chromatin immunoprecipitation (ChIP) and ChIP-Seq

B cells (1 × 10^7^) were treated with disuccinimidyl glutarate (1 mM) for 30 min and then washing with cold PBS before chromatin crosslinking by 1% formaldehyde in PBS at 25°C for 10 min. After quenching with 100 mM of glycine (pH 8.0) and washing with cold PBS containing a cocktail of protease inhibitors (Sigma-Aldrich), B cells were resuspended in SDS-lysis buffer (20 mM Tris-HCl, pH 8.0, 200 mM NaCl, 2 mM EDTA, 0.1% (w/v) sodium deoxycholate, 0.1% (w/v) SDS, and protease inhibitor cocktail). Chromatin was sonicated to yield approximately 200–600-bp DNA fragments, pre-cleared with agarose beads conjugated with protein A (ThermoFisher) and then incubated with polyclonal rabbit Ab to BATF3 or NF-kB p65. After overnight incubation, immune complexes were isolated using agarose-beads conjugated with protein A, washed and eluted in buffer containing 50 mM Tris-HCl, 0.5% SDS, 200 mM NaCl and 100 μg/ml Proteinase K (pH 8.0). Eluates were incubated at 65°C for 4 h to reverse formaldehyde crosslinks. DNA was purified using a QIAquick PCR purification column (Qiagen).

For ChIP-Seq, after the size and quality check on an Agilent Bioanalyzer 2100, DNA samples were used to prepare the library for sequencing at a HiSeq 3000. As a control, input DNA was reverse crosslinked and prepared using the same library preparation kit. After mapping of sequencing reads to the mouse mm10 genome and removal of duplicated reads using bowtie2, peak calling was performed by MACS2 using default settings except not building the shifting model and set extension size 200bp. Bigwig files were generated for visualization in the Integrative Genomics Viewer (IGV 2.3.81). Reads in peaks were counted by annotatePeaks.pl (–d option) from HOMER.

### Bioinformatics

To investigate biological pathways associated with DEGs in the core transcriptional signature of B cells stimulated *in vitro* with selected conditions, the ingenuity pathway analysis (IPA, Qiagen) was performed using aggregate mRNA profile was analyzed to identify statistically overrepresented pathways based on FDR-adjusted p values <0.05 and to infer pathway activation status based on Z scores. Alternatively, DEGs were manually curated and compared to multiple public databases, including Gene Ontology (GO) and Kyoto Encyclopedia of Genes and Genomes (KEGG), for enrichment analysis. DEGs were also directly compared to curated gene signatures to identify statistically significant concordance in the expression using the gene set enrichment analysis (GSEA) algorithm. To identify from 1500 curated transcription factors those that were inducible by the same IL-27-inducing stimuli and mediated IL-27 induction, expression of 411 genes with at least 5 counts per million reads in any of the 14 samples of *in vitro* stimulated B cells and that of *Il27p28* were subjected to unsupervised pair-wise correlation analysis using the *cor* function in R, yielding a matrix of correlation coefficients, which were visualized by a heatmap generated using the ggplot2 package in R. For scRNA-Seq, DEGs were subjected to principle component analysis (PCA) to identify subsets.

For ChIP-Seq, analysis of enriched motifs was performed with HOMER to identify *bona fide* NF-kB p65 target sites as well as *cis-*elements that were close to such target sites and occupied by a different transcription factor(s), with which NF-kB interacted with. Known motif enrichment in the peak sets for each stimulus was compared against the whole genome, whereas enrichment in the different differentially bound (DB) groups was compared against all the NF-kB p65 peaks. Peak location comparison between conditions was done with the mergePeaks command from HOMER using –d 500 setting and -venn option. To identify NF-kB binding to the promoter region of induced B cells, the BETA algorithm was used to integrate the ChIP-Seq and RNA-Seq datasets and return a score of interested genes based on their distance to TSSs (based on ChIP-Seq) and RNA-Seq expression profile.

### Immunoblotting

B cells (2×10^6^) were resuspended in lysis buffer supplemented with Halt™ Protease & Phosphatase Inhibitors Cocktail (ThermoScientific). After sonication, cell lysates were separated by SDS-PAGE and transferred onto PVDF membranes for immunoblotting. Membranes were then stripped with Restore™ PLUS Western Blot Stripping Buffer (ThermoScientific) for re-blotting.

### Histopathology

VV_WR_-infected mice were sacrificed at the pre-determined endpoint by cervical dislocation and infused with 2ml of 4% paraformaldehyde through intra-tracheal injection. The fixed lung tissues were harvested and then paraffin embedded for staining of tissue sections (5 μm) by hematoxylin and eosin staining. Lung pathology was evaluated by two independent investigators blinded to the identity to examine: 1. alveolar septa widened by swollen mononuclear cells which contain granular eosinophilic cytoplasm; 2. necrosis of cells within alveolar septa, karyorrhexis and mild infiltration of alveolar walls by neutrophils, fibrin; 3. exudation of leukocytes into alveolar air spaces; 4. edema (eosinophilic proteinaceous material within distal air sacs (intra alveolar edema) sometimes intermixed with aggregates of red cells (hemorrhage); 5. aggregates of mononuclear cells (lymphocytes and mononuclear/macrophage) in peribronchiolar and perivascular spaces; 6. the mucosa of bronchioles hyperplastic (pilling up) or extended (type II pneumocyte hyperplasia); desquamated or necrotic epithelial cells occasionally present in the lumens; 7. fibrosis; 8. blood vessel changes, including perivascular and interstitial edema, thrombosis and margination of neutrophils.

### Statistical analysis

All statistical analyses were performed using Excel (Microsoft), GraphPad Prism^®^ or R software environment. Differences in RNA transcript expression were determined by Deseq2, which estimate variance-mean dependence in count data and test for differential expression based on a model using the negative binomial distribution. Spearman correlation analysis was used to measure the strength and direction of gene expression correlations. The student *t-*test was used to analyze the group differences, assuming equal variances. The Mantel–Cox test was used for survival analysis.

### Reagents

All antibodies are listed in **Table S19**, and the sequence of all primers are listed in **Table S20**.

---

**fig. S1.**
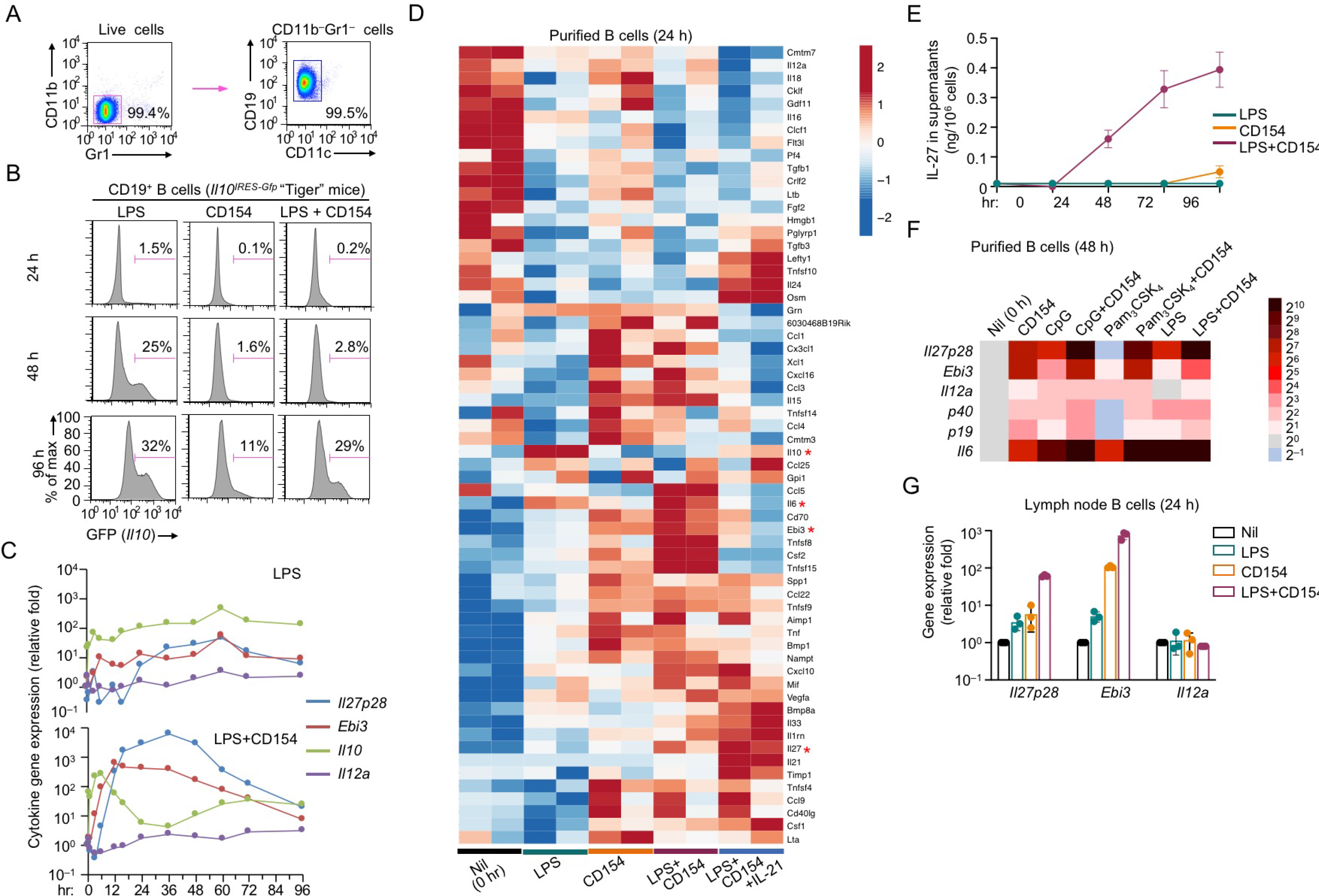
Specific induction of IL-27 in B cells by TLR ligands plus CD154. **(A)** Purity of B cells after negative selection and used for *in vitro* stimulation **(B)** FACS analysis of GFP induction in *Il10^IRES-Gfp^* B cells (from *Il10^tm1Flv^* mice, JAX #008379) Representative of two independent experiments. **(C)** qRT-PCR analysis of cytokine gene transcript levels in purified speen B cells after stimulation at time points as indicated. Data were normalized to *Cd79b* expression and expressed as the ratio to values of freshly isolated B cells (0 h). Representative of two independent experiments. **(D)** Heatmap of expression of genes with cytokine and chemokine activities (JAXInformatics) differentially expressed in B cells after stimulation, as indicated, for 24 h. Data are from two independent RNA-Seq experiments. Only genes that were differentially expressed as compared to those in freshly isolated B cells (nil. h 0) and showed >= 50 reads in at least one sample were depicted. **(E)** ELISA of IL-27 secreted into the supernatants of purified spleen B cells after stimulation, as indicated (mean and s.d., triplicates). **(F)** qRT-PCR analysis of cytokine transcript levels in purified spleen B cells, as induced by indicated stimuli after 48 h of stimulation. Data were normalized to *Cd79b* expression and expressed as the ratio to values of freshly isolated B cells (0 h). Representative of two independent experiments. **(G)** qRT-PCR analysis of gene transcript levels in purified lymph node B cells 48 h after stimulation. Data were normalized to *Cd79b* expression and expressed as the ratio to values of freshly isolated B cells. Representative of two independent experiments.

**fig. S2.**
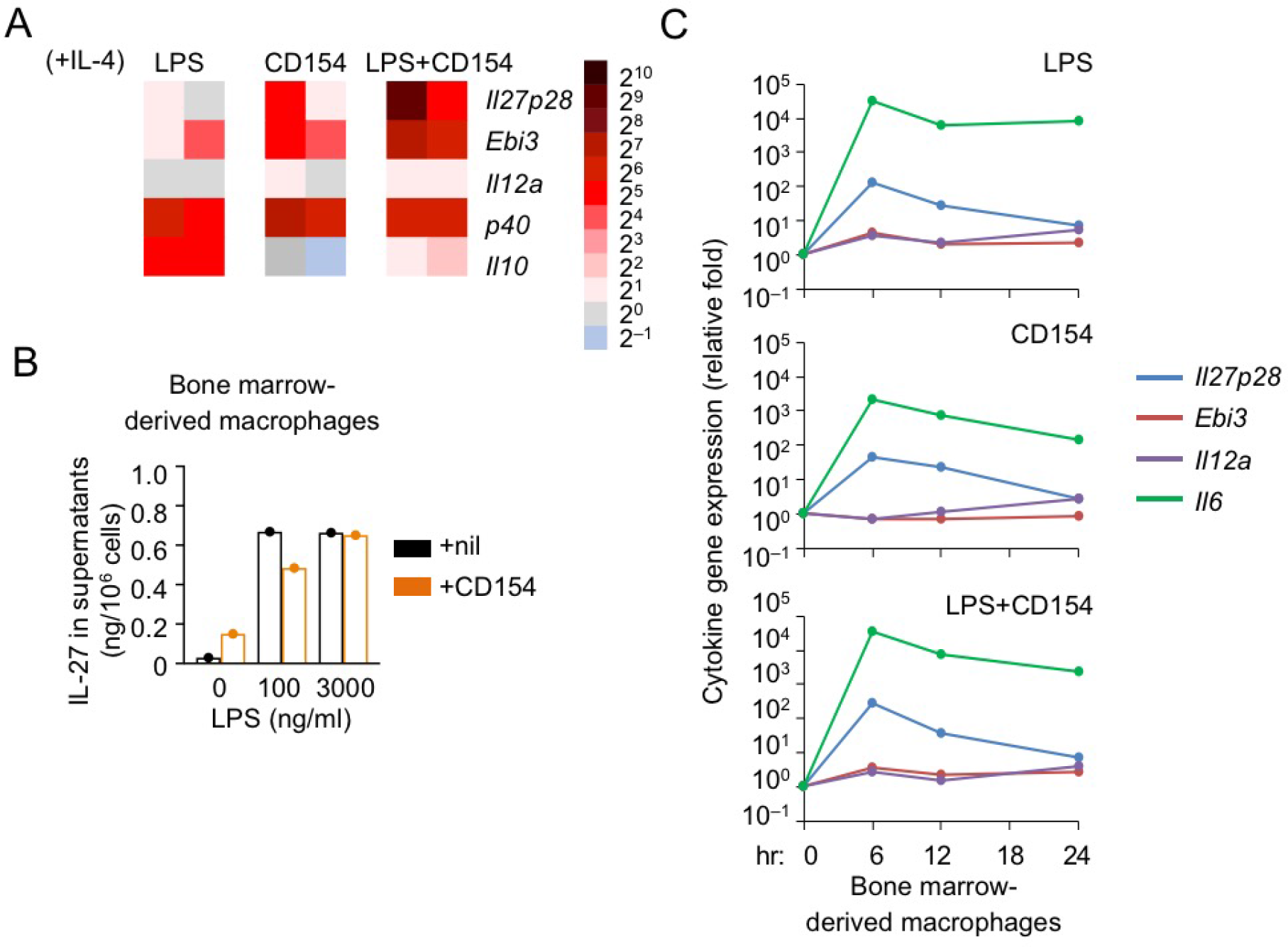
IL-27 induction in B cells and macrophages. **(A)** qRT-PCR analysis of gene induction in purified spleen B cells after stimulation, as indicated, for at 24 or 48 h. Data were normalized to *Cd79b* expression and expressed as the ratio to values of freshly isolated B cells (0 h). Representative of two independent experiments. **(B)** ELISA of IL-27 secreted into the supernatants of bone marrow-derived macrophages after stimulation, as indicated, for 48 h. Representative of two independent experiments. **(C)** qRT-PCR analysis of cytokine transcript levels, as indicated, in bone marrow-derived macrophages, as induced by indicated stimuli after 48 h of stimulation. Data were normalized to *Gapdh* expression and expressed as the ratio to values of non-stimulated macrophages. Representative of two independent experiments.

**fig. S3.**
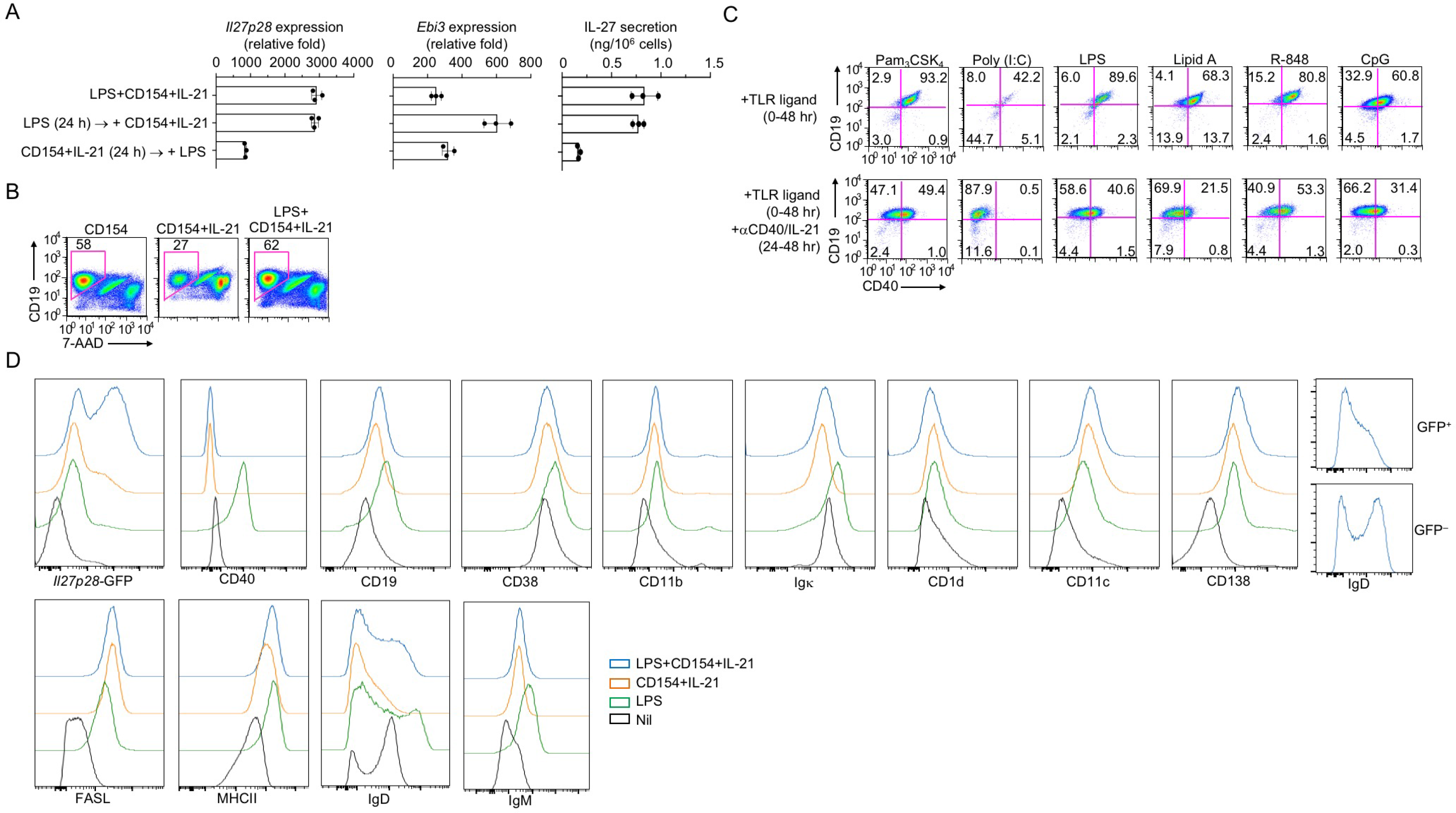
Induction of IL-27 in B cells by sequential stimulation of LPS and CD154. **(A)** qRT-PCR analysis of *Il27p28* and *Ebi3* transcript levels (left two panels) and ELISA of IL-27 secreted into the supernatants (right panel) of purified spleen B cells after co-stimulation with LPS, CD154 or IL-21, or sequential stimulation, as indicated (mean and s.d., triplicates). **(B)** Flow cytometry analysis of cell viability (7-AAD^-^) in B cells stimulated, as indicated, for 48 h. Representative of five independent experiments. **(C)** Flow cytometry analysis of CD40 surface expression in B cells stimulated, as indicated, for 24 h. **(D)** Flow cytometry analysis of expression of CD40 and selected surface markers in B cell stimulated, as indicated, for 48 h. Representative of two independent experiments.

**fig. S4.**
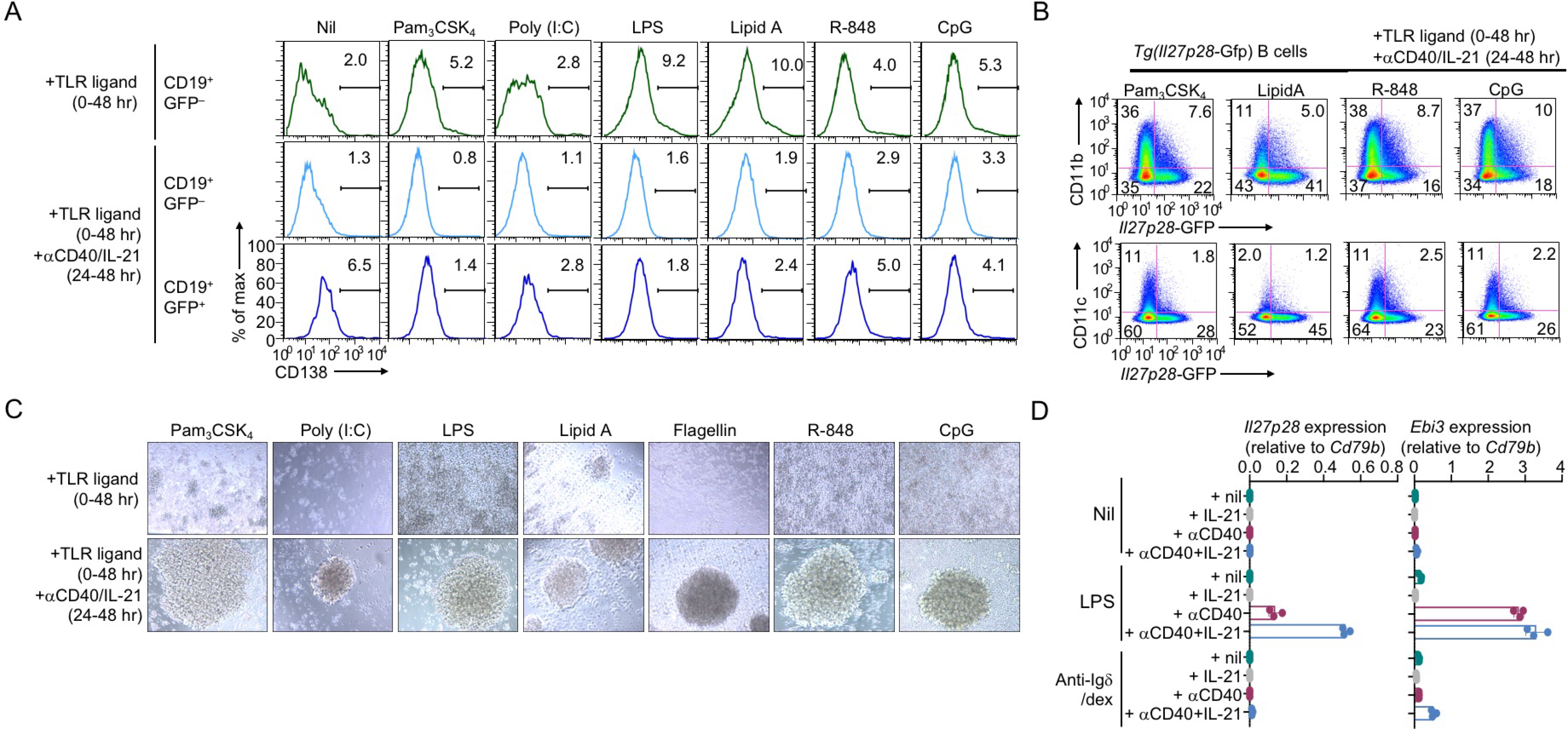
Specific induction of IL-27 in B cells by TLR priming and exposure to αCD40 and IL-21. **(A,B)** Flow cytometry analysis of CD138 **(A),** CD11b and CD11c **(B)** expression in purified *Tg*(*I112p28-Gfp*) B cells after priming with a TLR ligand, as indicated, for 24 h and then added αCD40 and /IL-21 for 24 h. Representative of three independent experiments. **(C)** Examination of purified B cells after priming with a TLR ligand, as indicated, for 24 h and then added αCD40 and /IL-21 for 24 h. Representative of six independent experiments. **(D)** qRT-PCR analysis of *Il27p28* and *Ebi3* transcript levels in purified spleen B cells after priming with nil, LPS or anti-lgδ/dex for 24 h and then added αCD40 and/or IL-21 for 24 h. Data were normalized to *Cd79b* expression and expressed as the ratio to values of freshly isolated B cells (0 h).

**fig. S5.**
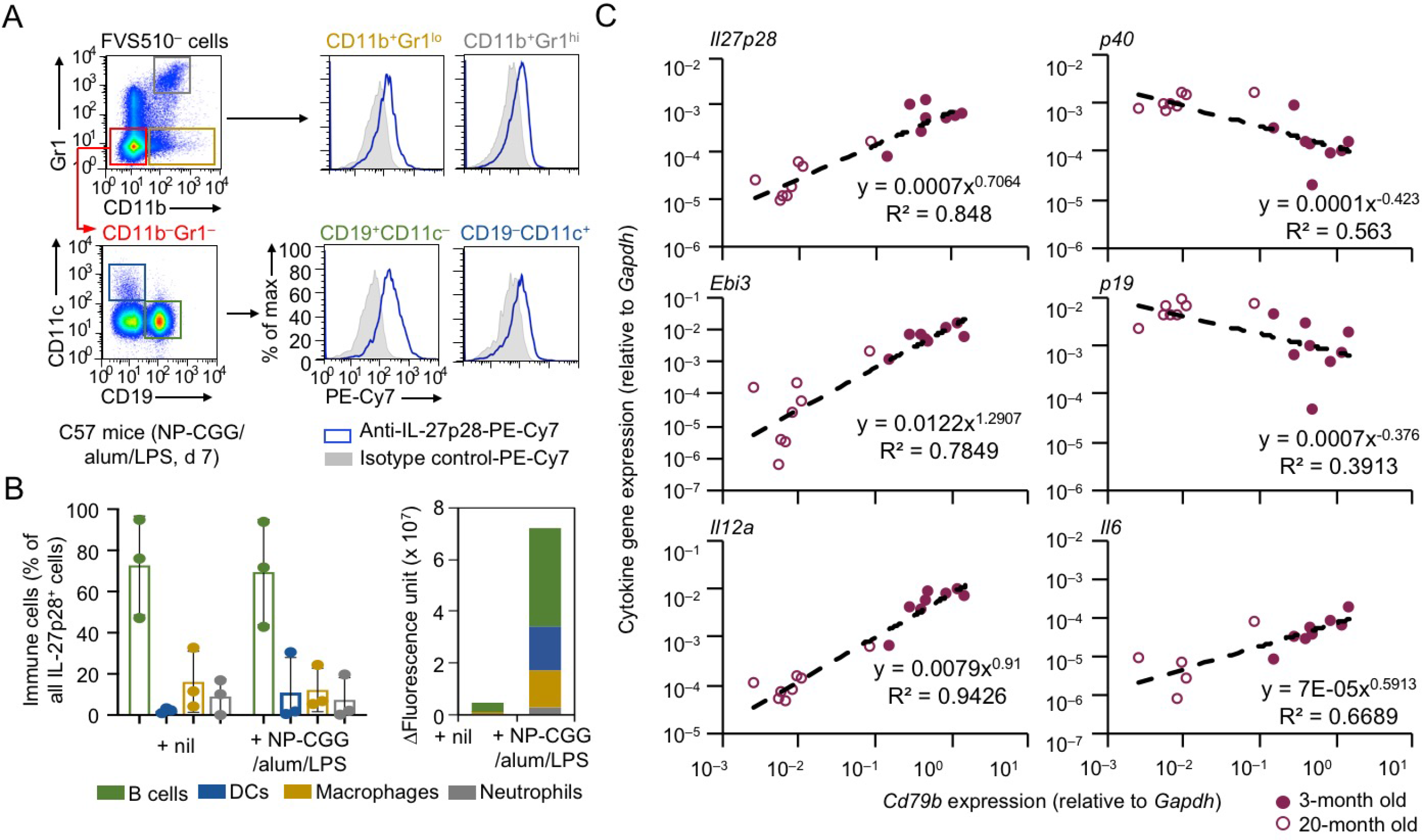
Expression of IL-27 by B cells *in vivo.* **(A)** Intracellular staining with a IL-27p28-specific Ab and isotype-matched IgG control Ab and flow cytometry analysis of the fluorescence signals in B cells (CD11b-Gr1^-^CD19^+^CD11c^-^), DCs (CD11b^-^Gr1^-^CD19^-^CD11c^+^), macrophages (CD11b^+^Gr1^lo^) and neutrophils (CD11b^+^Gr1^hi^) in C57 mice 7 d after immunization with NP-CGG, alum and LPS. Representative of three independent experiments. **(B)** Proportions of different immune cell populations, as defined in (A), in IL-27p28+ cells, as analyzed by flow cytometry, in C57 mice before or 7 d after immunization (left panel); also depicted is the cumulative fluorescence unit of each population, as calculated by multiplication of the number of cells with the subtraction of fluorescence arising from isotype-matched IgG Ab staining from that of anti-IL-27p28 staining (right panel). **(C)** Correlation analysis of transcript levels of cytokine-encoding genes, as indicated, with that of *Cd79b*, as determined by qRT-PCR, in 8 young (3-month old) and 8 aging (20-month old) mice that were not immunized. Data were normalized to *Gapdh* expression for the calculation of power regression and R^2^ value (Excel).

**fig. S6.**
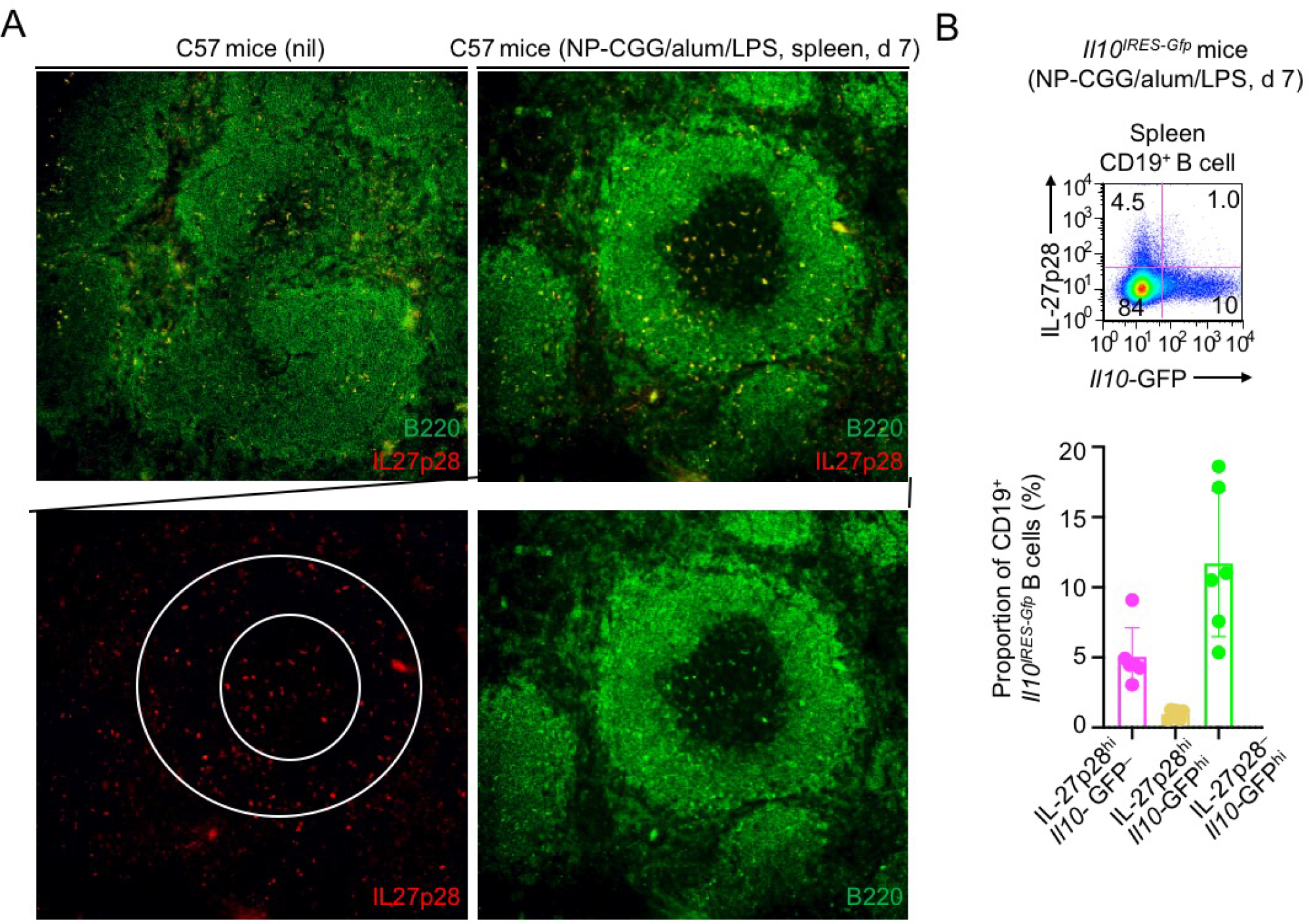
IL-27-producing B cells in mice upon immunization. **(A)** Immunofluorescence microscopy of IL-27p28^+^B220_+_ cells in the spleen of C57 mice after immunization. A representative follicle in the filed is shown. **(B)** Flow cytometry analysis of IL-27p28 and GFP (indicating *Il10* transcription) in B cells in immunized *Il10^IRES-Gfp^* mice. Data quantification (lower panel) are from three independent experiments.

**fig. S7.**
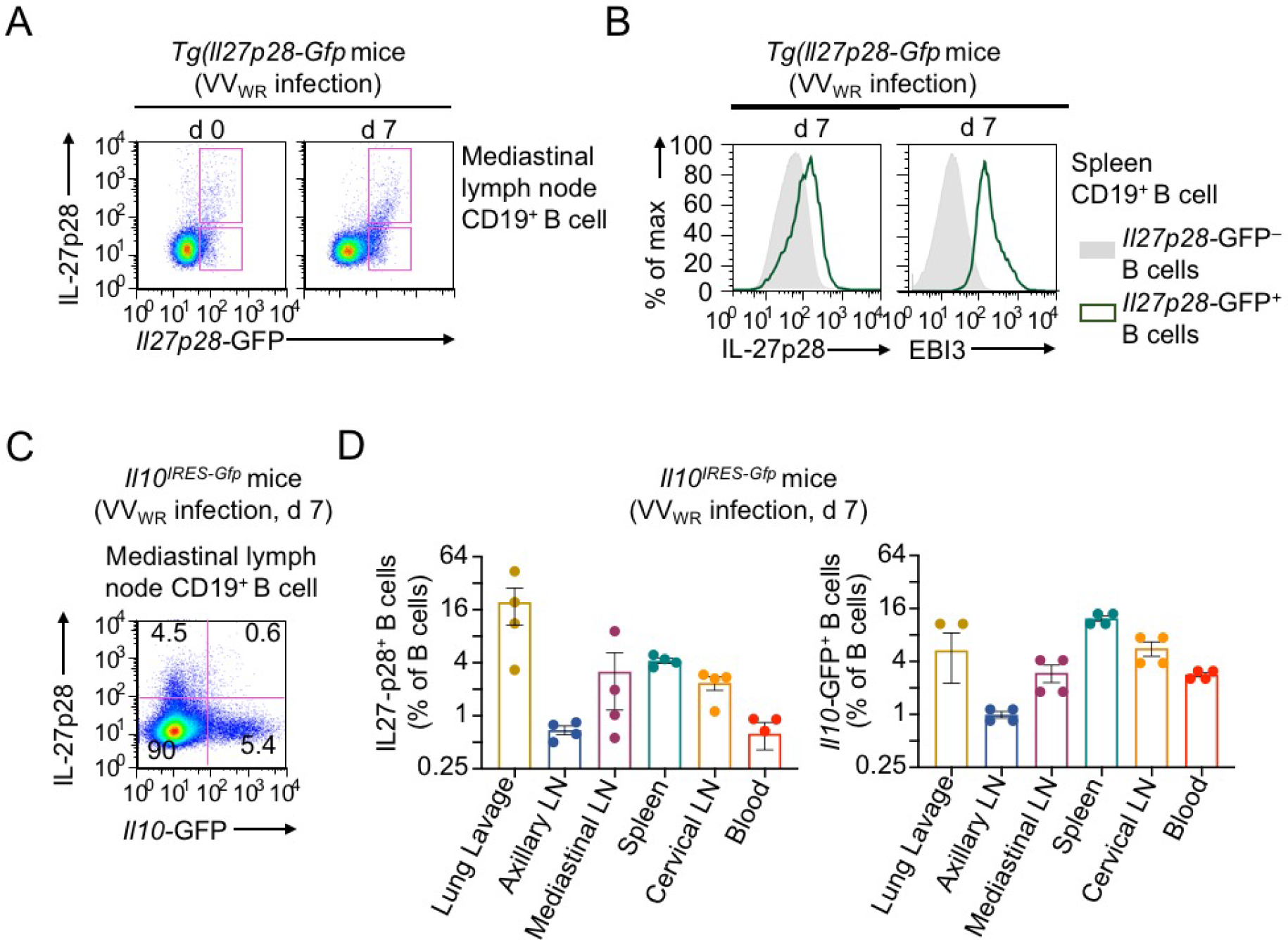
IL-27-producing B cells are induced in mice upon VV_WR_ infection. **(A)** Flow cytometry analysis of IL-27p28 and GFP (indicating *Il27p28* transcription) expression to quantify IL-27p28 induction in mediastinal lymph node B cells *Tg(Il27p28-Gfp*) mice before and after infection with VV_WR_ infected. **(B)** Flow cytometryanalysis of IL-27p28 and EBI3 in infected *Tg(Il27p28-Gfp*) mice. Representative of three independent experiments. **(C,D)** Flow cytometry analysis of IL-27p28 and GFP (indicating *Il10* transcription) expression (**C**, representative flow cytometry plot) to quantify IL-27p28 protein and *Il10* transcription induction in infected *Il10^IREs-Gfp^* mice (**D**, and quantification of data from different tissues and organs).

**fig. S8.**
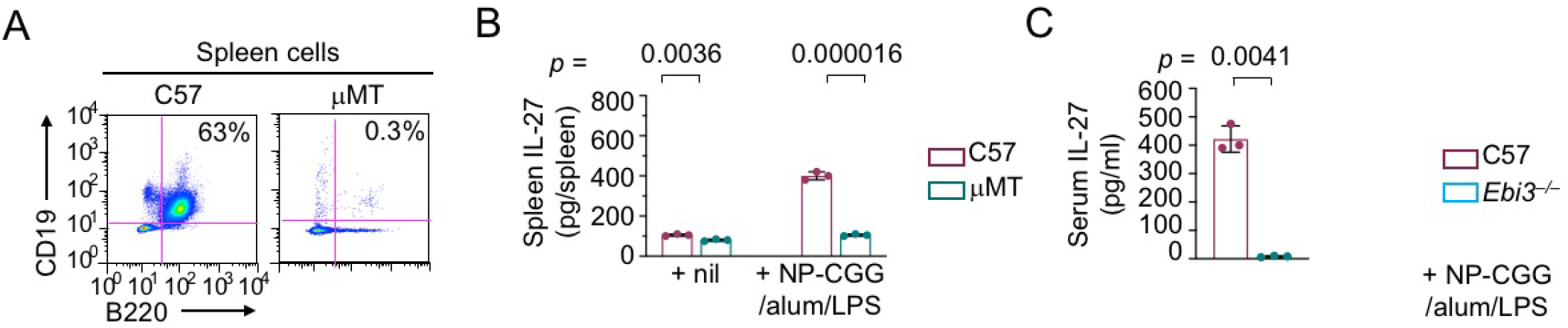
Reduced IL-27 production in mice lacking B cells. **(A)** Flow cytometry analysis of lack of B cells in μMT mice. **(B)** ELISA of IL-27 in the spleen in C57 and μMT mice before and after (d 7) of immunization. **(C)** ELISA of IL-27 in the circulation of C57 and *Ebi3^−/−^* mice after immunization for 7 d.

**fig. S9.**
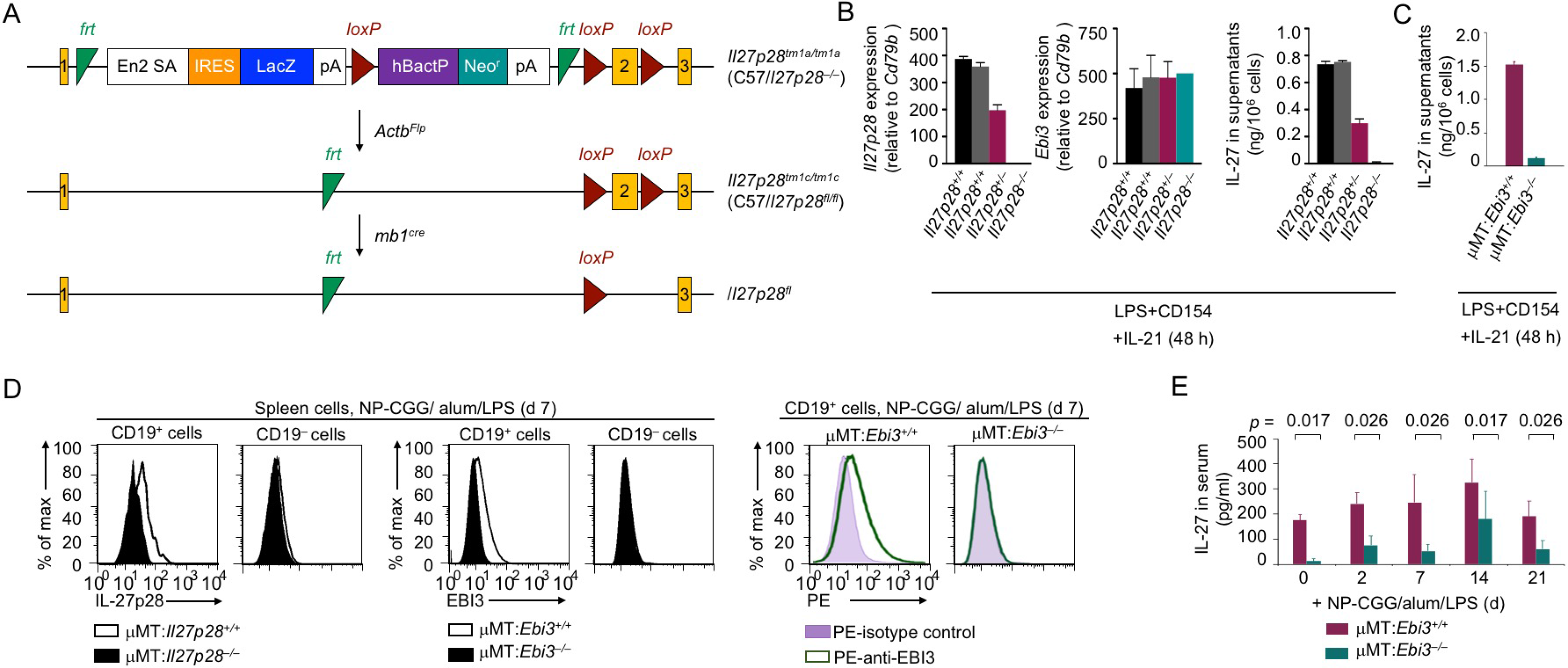
B cell-specific deficiency in *Il27p28* or *Ebi3* results in decreased IL-27 production. **(A)** Schematics of *Il27p28^-^* allele, due to the interference of *Il27p28* transcription from a large insert between exon 1 and exon 2 in the *Il27p28^,m1a^* allele, and the generation of *Il27p28^fl^* allele, which is independent from that used in this study. **(B)** qRT-PCR analysis (left) and ELISA (right) for the confirmation of lack of *Il27p28* transcription and IL-27 secretion in *Il27p28^−/−^* B cells stimulated *in vitro.* qRT-PCR Data were normalized to *Cd79b* expression and expressed as the ratio to values of freshly isolated B cells (0 h). **(C)** ELISA for the confirmation of lack of IL-27 secretion in μMT:*Ebi3^−/−^* B cells stimulated *in vitro.* **(D)** Intracellular staining and flow cytometry analysis for the confirmation of lack of IL-27p28 and EBI3 expression in immunized μMT:*Il27p28^−/−^* and μMT:*Ebi3^−/−^* mice, respectively. Representative of two independent experiments. **(E)** ELISA of reduced IL-27 in the circulation in μMT:*Ebi3^−/−^* mice at different timepoints after immunization.

**fig. S10.**
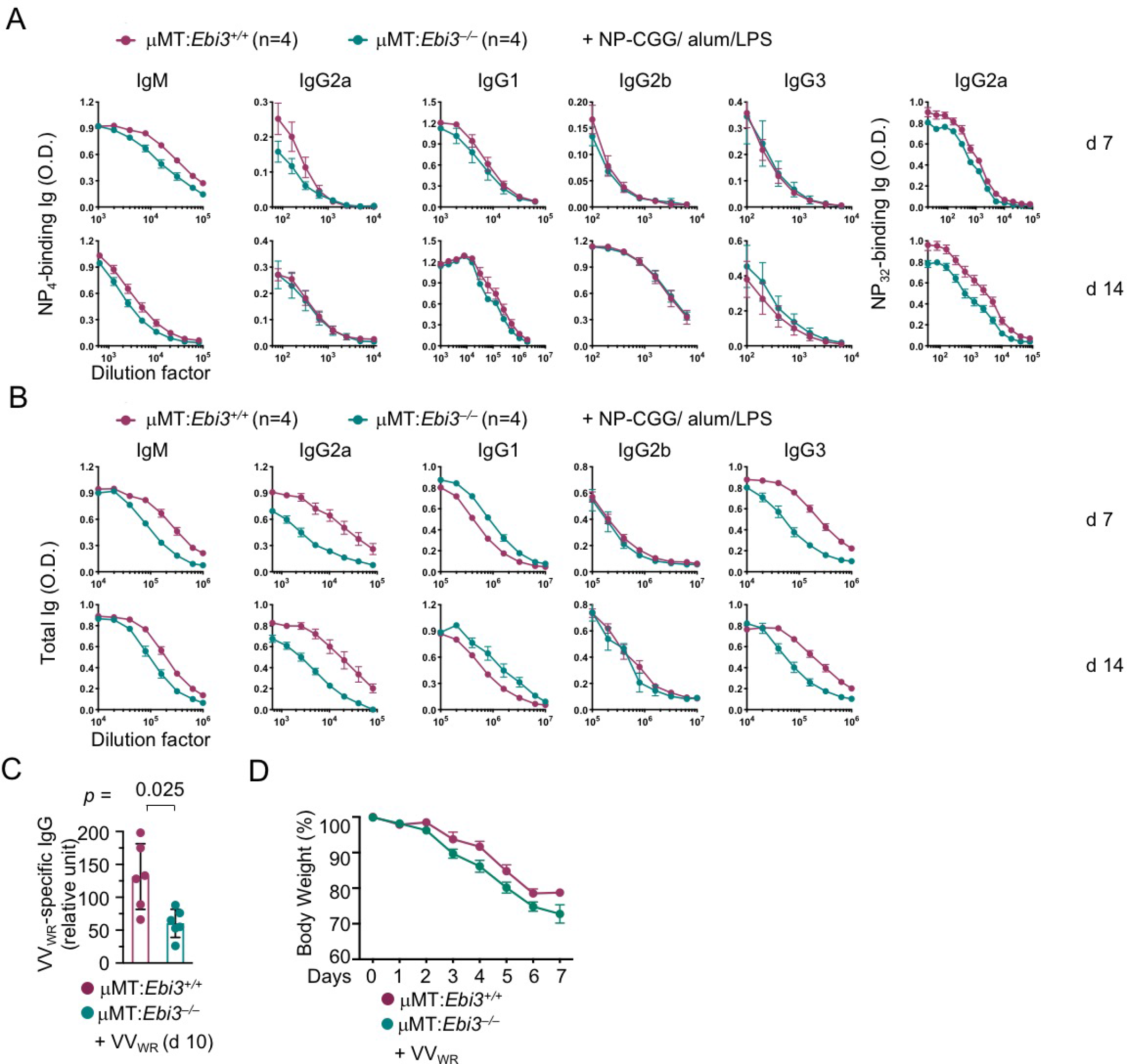
Defective IgG2a responses in μMT:*Ebi3^−/−^* mice. **(A)** ELISA of NP_4_-binding (high-affinity) and (all specific) NP_32_-binding Ig isotypes in immunized μMT:*Ebi3^−/−^* mice. **(B)** ELISA of total Abs of different Ig isotypes in immunized VV_WR_-infected μMT:*Ebi3^−/−^* mice, as in **(A). (C,D)** ELISA of virus-specific total IgG in infected μMT:*Ebi3^−/−^* mice **(C)** and the loss of body weight in such mice **(D)**.

**fig. S11.**
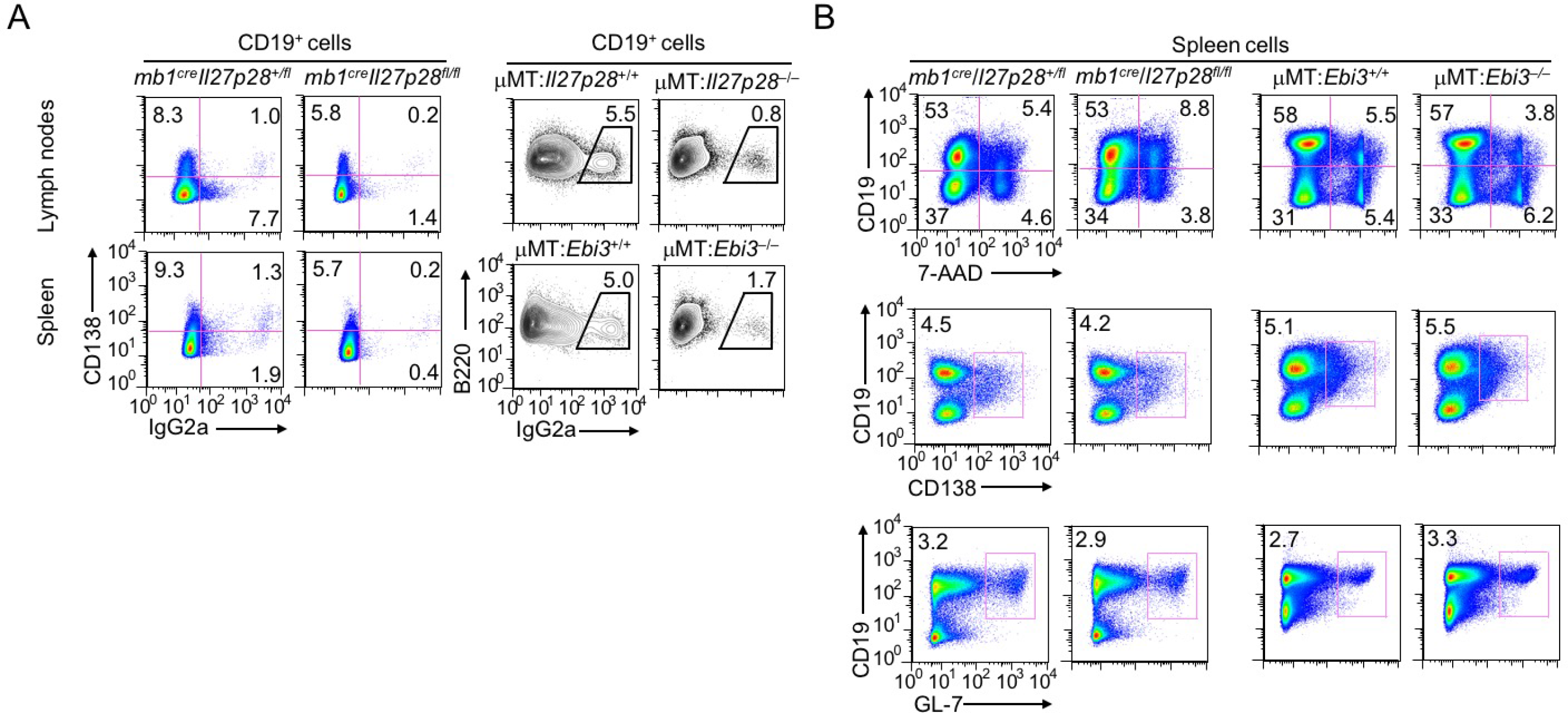
Impairment in CSR to IgG2a in mice with B cell-specific deficiency in *Il27p28* or *Ebi3.* **(A)** Flow cytometry analysis of IgG2a and/or CD138 in spleen and lymph node B cells in immunized *mb1^cre^Il27p28^fl/fl^*, μMT:*Il27p28^−/−^* and μMT:*Ebi3^−/−^* mice mice, as well as their respective wildtype mouse counterparts. Representative of two independent experiments. **(B)** Flow cytometry analysis of cell viability, CD138 and GL-7 in immunized mice, as in **(A)**. Representative of three independent experiments.

**fig. S12.**
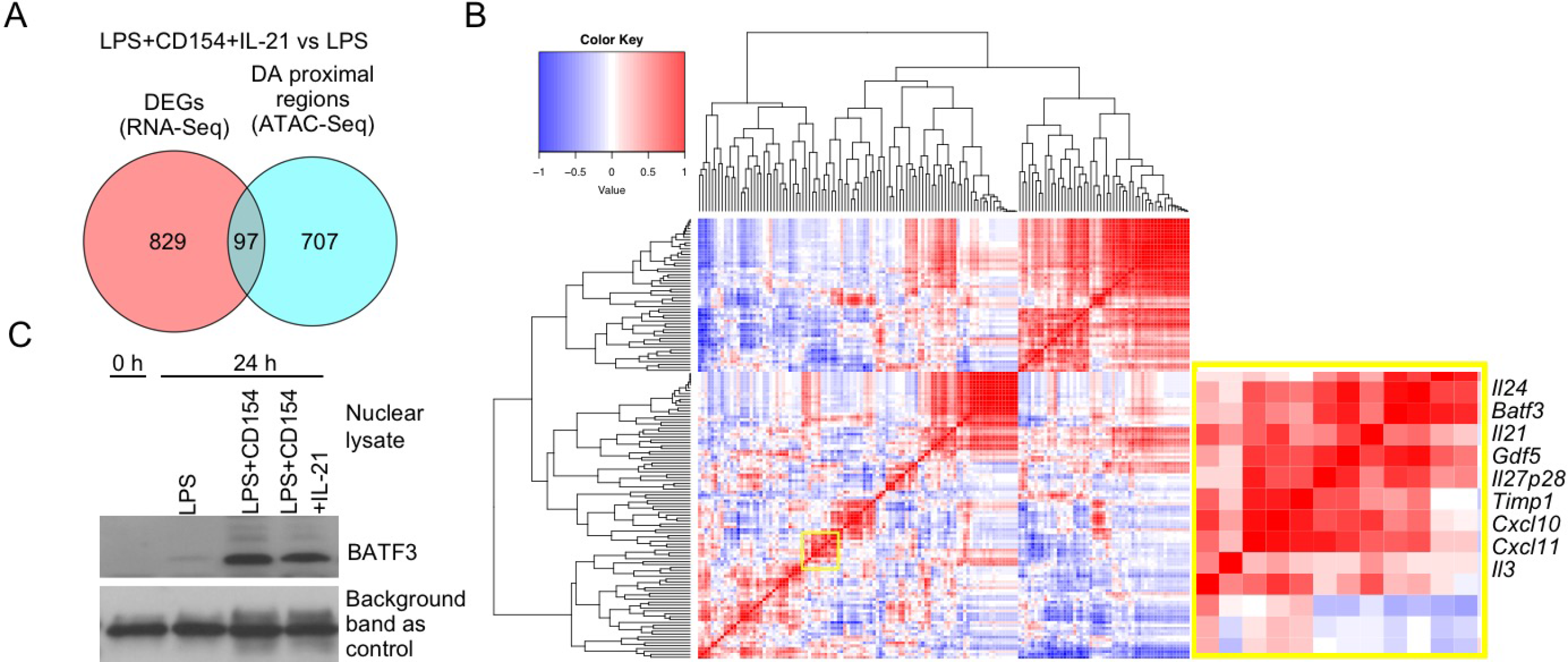
BATF3 is upregulated in B cells induced to express IL-27 *in vitro.* **(A)** Venn diagram of 97 genes that had DARs and DEGs in B cells stimulated with LPS plus CD154 and IL-21 to induce IL-27, as compared to LPS-stimulated B cells. *Il27p28* is one of the 97 genes. **(B)** Correlation analysis of the expression level of of *Batf3* and that of 120 expressed cytokine and chemokine-encoding genes in duplicate B cells stimulated *in vitro* with nil (0 h), (CSR-inducing) LPS, CD154 and CD154 plus IL-4, as well as (IL-27-inducing) LPS plus CD154, CD154 and IL-21, and LPS plus CD154 and IL-4, for 24 h. **(C)** Immunoblotting of induced BATF3 in the nucleus of B cells stimulated with LPS plus CD154 and/or IL-21. Representative of two independent experiments.

**fig. S13.**
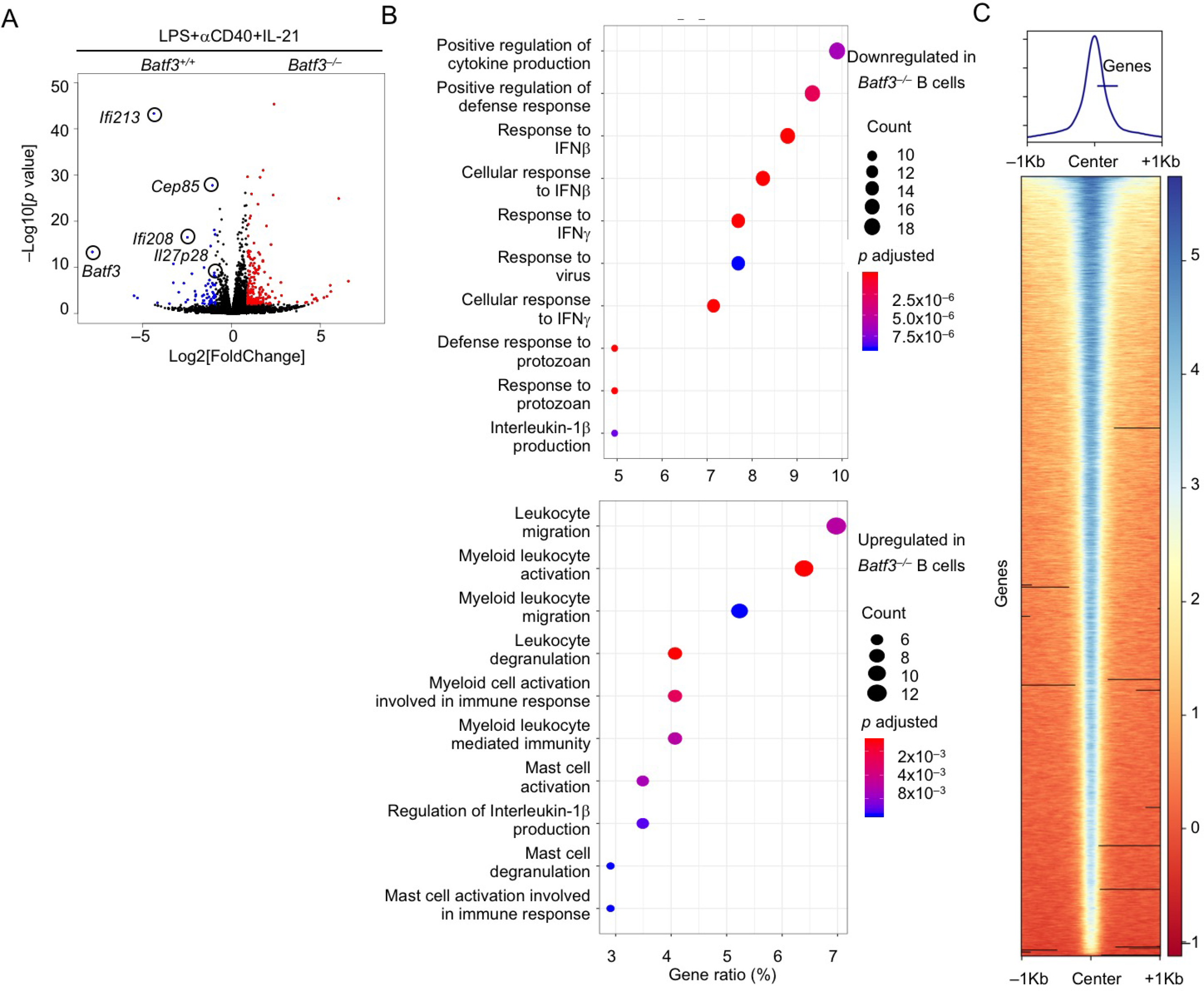
BATF3 mediates IL-27 induction in B cells *in vitro.* **(A)** Volcano plot DEGs in *Batf3^−/−^* and *Batf3^+/+^* B cells after stimulation for 48 h. **(B)** GO analysis of DEGs in stimulated *Batf3^−/−^* and *Batf3^−/−^* B cells, as in **(A). (C)** Lack of significant DARs, as determined by ATAC-Seq, in *Batf3*^−/−^ and *Batf3^+/+^* B cells after stimulation with LPS plus αCD40 and IL-21 for 48 h.

**fig. S14.**
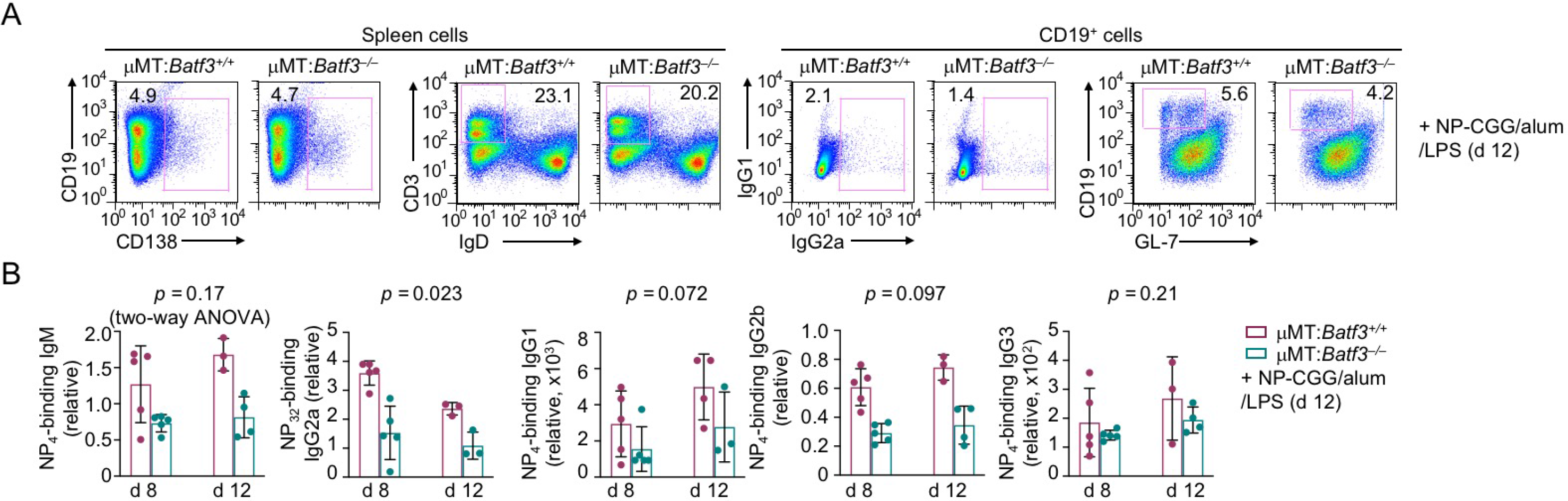
B cell-intrinsic role of BATF3 in mediating IL-27 induction and an antigen-specific IgG2a response. **(A)** Flow cytometry analysis of B and T cell proportions in the spleen (left) as well as B cell CSR and germinal center differentiation (right) in μMT:*Batf3^−/−^* mice and their wildtype counterparts immunized with NP-CGG plus alum and LPS for 12 d. Representative of three independent experiments. **(B)** ELISA of NP-specific Ig isotypes in μMT:*Baff3^−/−^* mice and their wildtype counterparts immunized with NP-CGG plus alum and LPS for 8 and 12 d.

**fig. S15.**
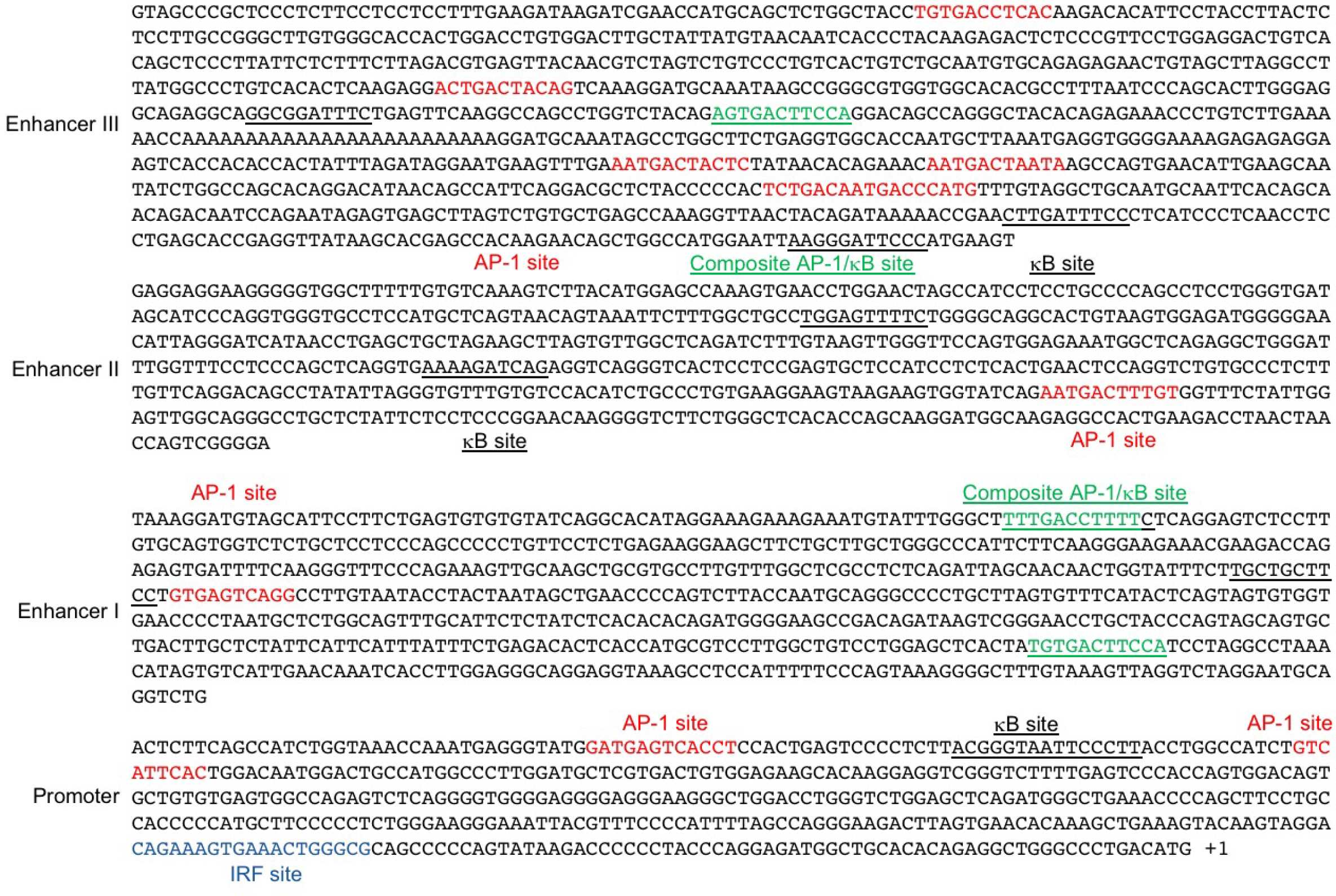
Putative binding sites in the promoter region and the three upstream enhancers in the *Il27p28* locus.

**fig. S16.**
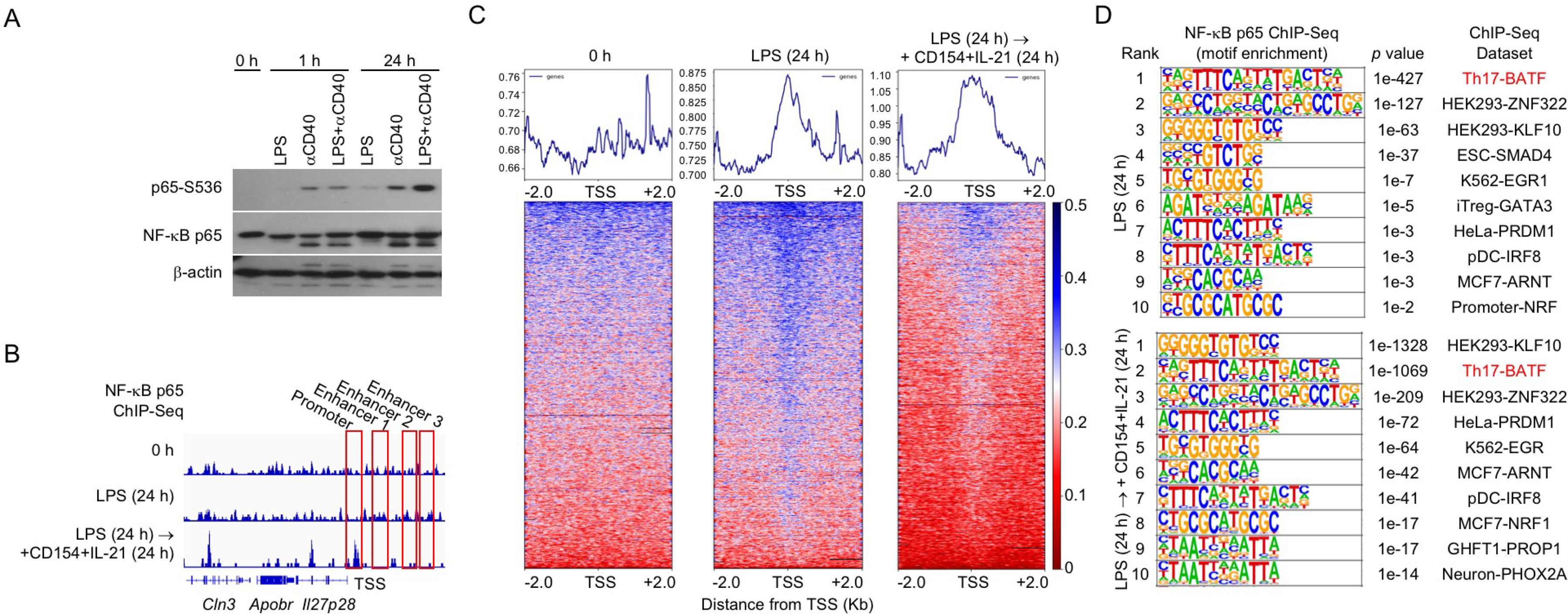
Genome-wide binding of NF-κB p65. **(A)** Immunoblotting of NF-κB p65 phosphorylation in B cells stimulated *in vitro*, as indicated. Representative of three independent experiments. **(B)** ChIP-Seq analysis of NF-κB p65-binding in the *Il27p28* locus in stimulated B cells, as indicated. **(C)** Heatmap of NF-κB p65 binding sites in the genome relative to the position of TSSs, as analyzed by ChIP-Seq. **(D)** Analysis of motifs enriched in NF-κB p65 binding sites.

**fig. S17.**
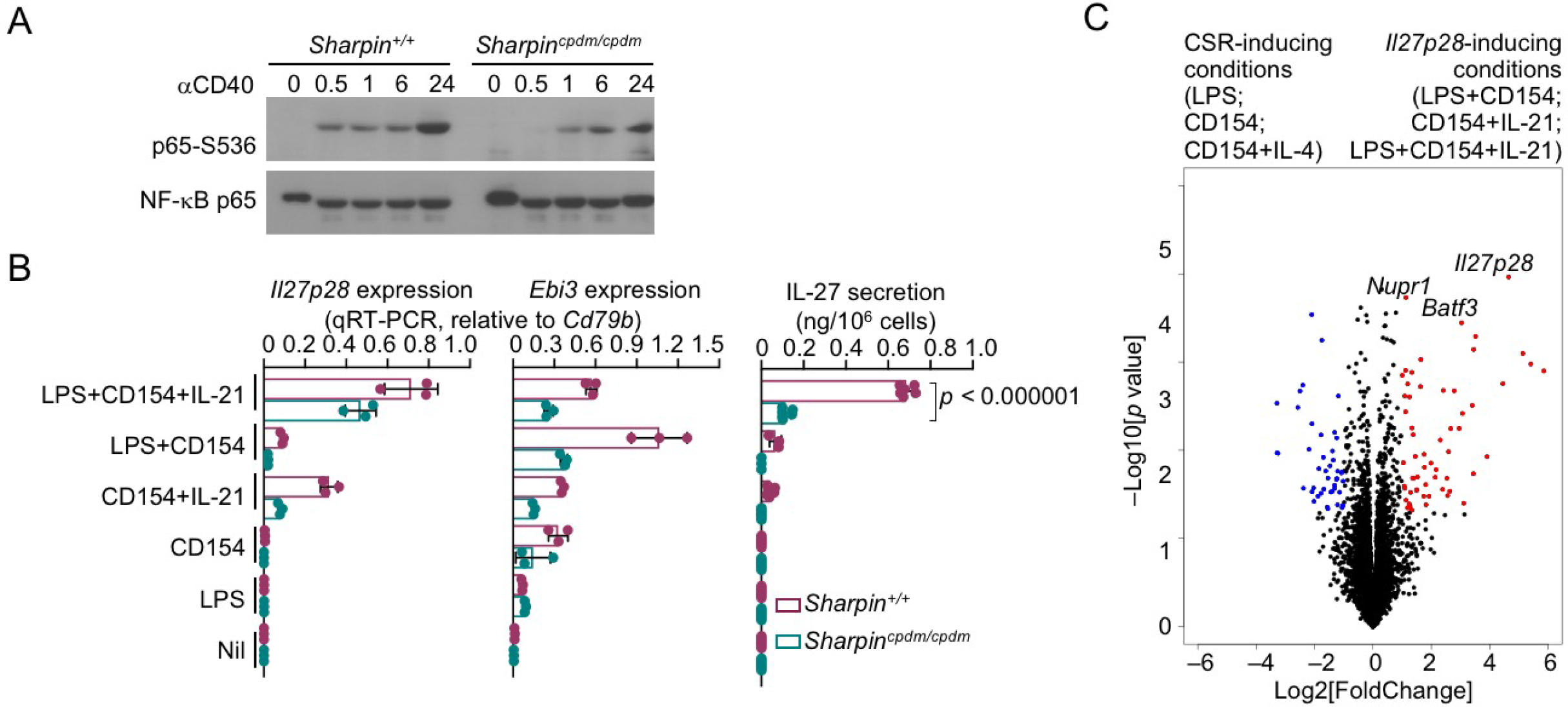
SHARPIN promotes NF-κB p65 activation and IL-27 induction. **(A)** Immunoblotting of phosphorylation of NF-κB p65 at Ser355 (p65-S355) in B cells stimulated *in vitro*, as indicated. Representative of two independent experiments. **(B)** qRT-PCR analysis of *Il27p28* and *Ebi3* expression in *Sharpin^+/+^* and *Sharpin^cpdm/cpdm^* B cells stimulated for 48 h (left and middle) and ELISA of IL-27 secretion by these cells (right). **(C)** Volcano plot depicting DEGs in B cells stimulated with *Il27p28*-inducing conditions and those stimulated with CSR-inducing conditions, as indicated, for 48 h.

**fig. S18.**
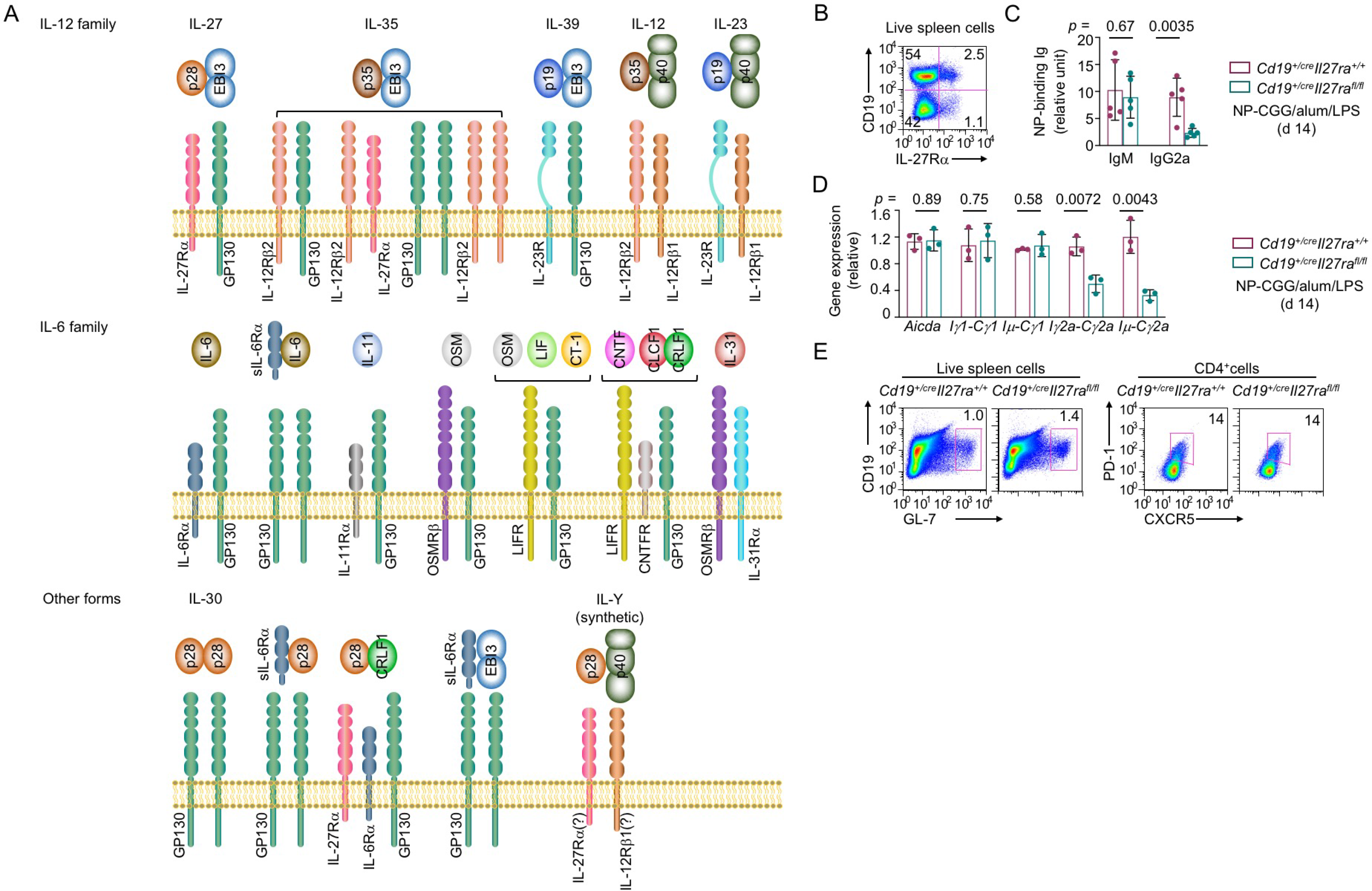
B cell-intrinsic role of IL-27 receptor signaling in IgG2a response *in vivo.* **(A)** Illustration of cytokines in the IL-12 family, IL-6 family and related forms as well as their respective receptors. **(B)** Flow cytometry analysis of IL-27Rα in spleen cells in non-immunized C57 mice. Representative of three independent experiments. **(C-E)** ELISA of NP-binding IgM and IgG2a **(C)**, qRT-PCR analysis of gene expression **(D)**, and flow cytometry analysis of B cells and T cells (E; representative of three independent experiments) in *Cd19^+/cre^Il27ra^fl/fl^* and *Cd19^+/cre^Il27^+/+^* mice immunized with NP-CGG plus alum with LPS for 14 d.

**fig. S19.**
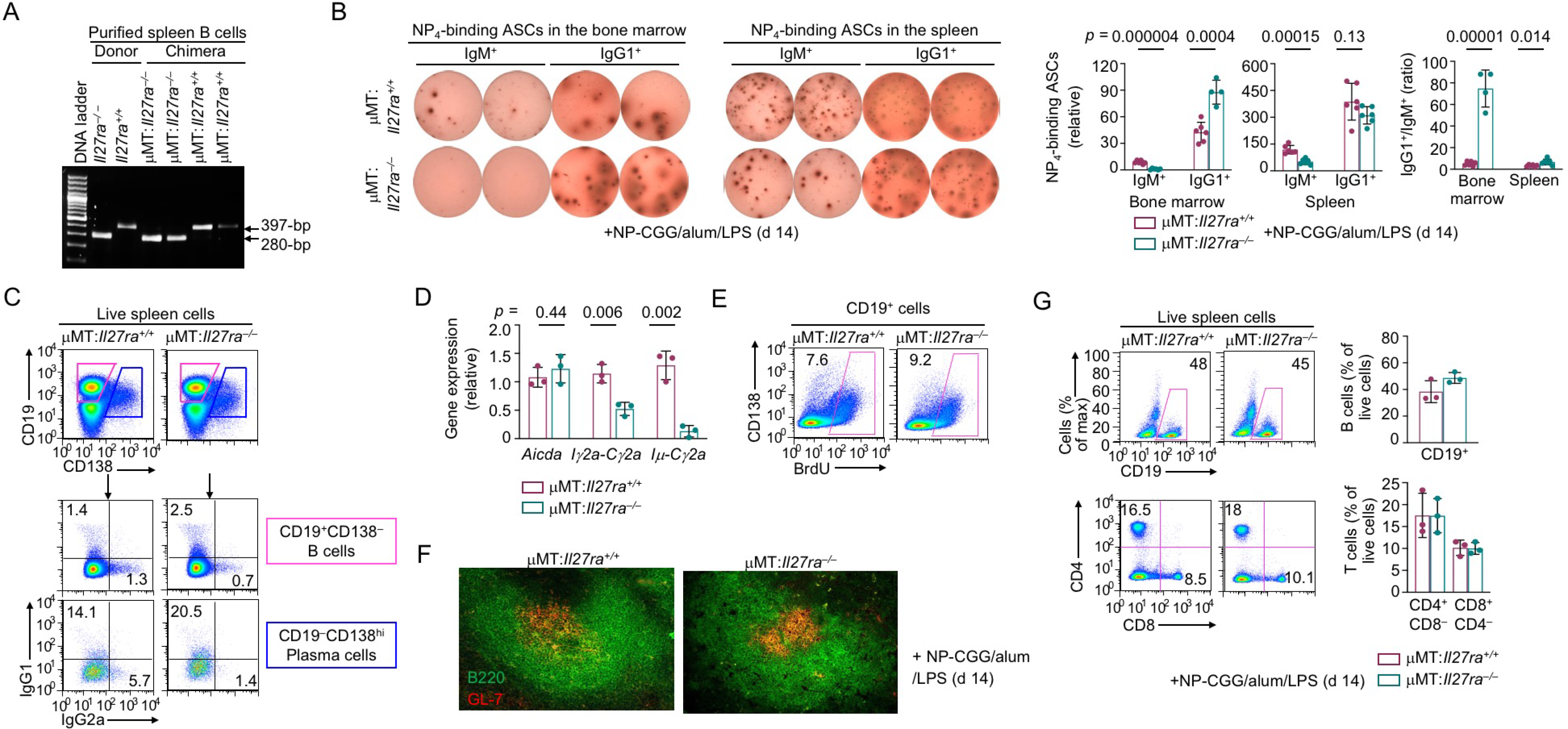
IL-27 receptor signaling mediates T-dependent IgG2a responses. **(A)** qPCR analysis of the B cell *Il27ra* genotype in mixed bone marrow chimera mice. **(B)** ELISPOT of ASCs secreting NP_4_-binding IgM and IgG1 in the bone marrow and spleen of immunized chimera mice. **(C)** Flow cytometry analysis of B cells and plasma cells, as defined by their surface CD19 and CD138 expression, and their expression of IgG2a and IgG1, as measured by intracellular staining in the spleen of immunized chimera mice (d 14). Representative of three independent experiments. **(D)** qRT-PCR analysis of CSR-related transcripts in spleen B cells isolated from immunized chimera mice (d 14). Data are normalized to *Cd79b* expression and expressed as the relative fold to values in one μMT:*Il27rα*^+/+^ mouse. **(E,F)** Flow cytometry and immunofluorescence microscopy analysis of B cell proliferation by BrdU incorporation **(E)** and germinal center differentiation **(F)** in immunized chimera mice (d 14). Representative of two independent experiments. **(G)** Flow cytometry analysis of the proportions of live B cells and T cells in immunized chimera mice (d 14).

**fig. S20.**
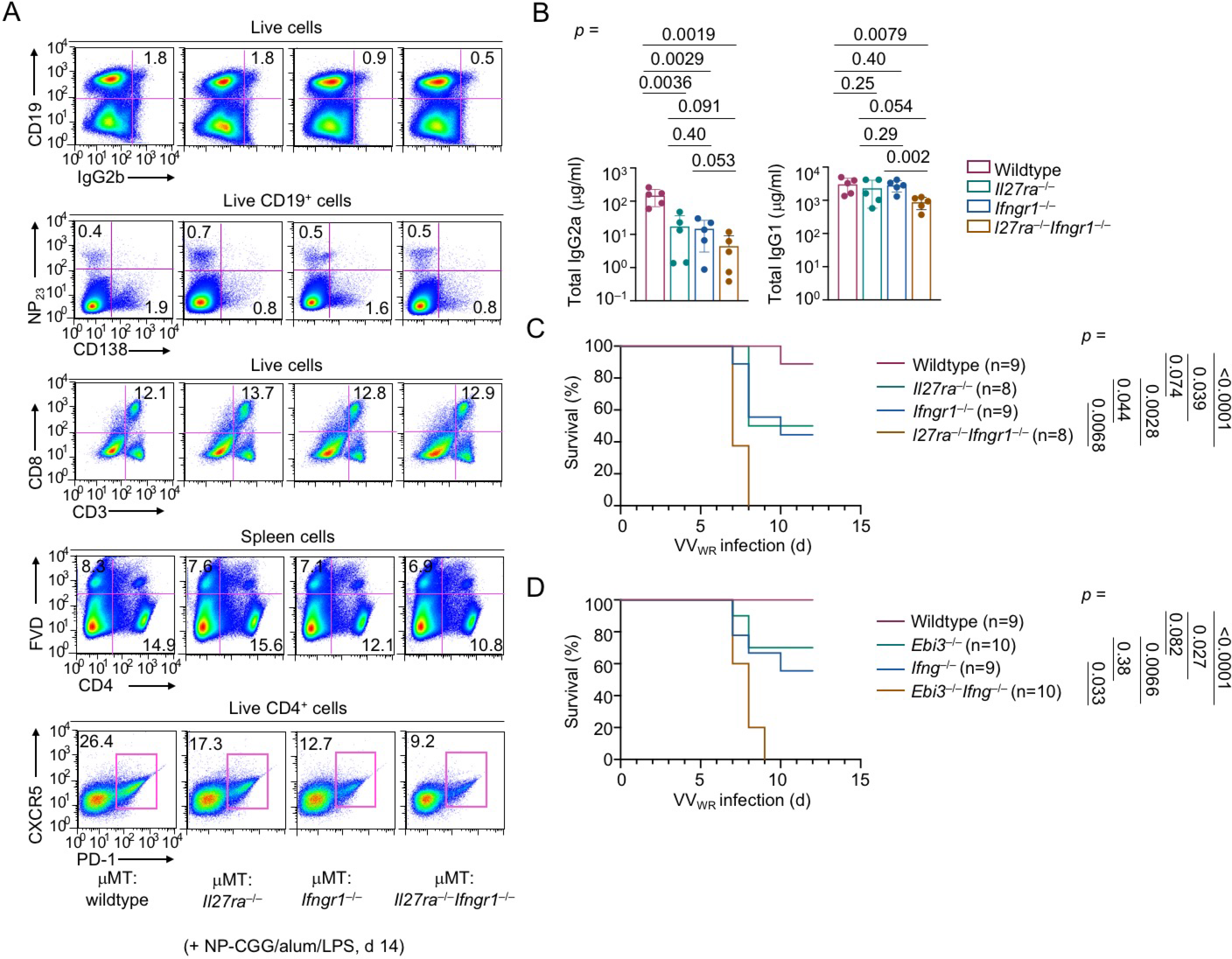
Synergy of IL-27 receptor and IFNγ receptor signals *in vivo.* **(A)** Flow cytometry analysis of B cells and T cells in double and single KO chimera mice, as indicated, after immunization. Representative of five mice in each group. **(B)** ELISA of total IgG2a and IgG1 in non-immunized mice with double and single KO in cytokine receptor genes, as indicated. **(C,D)** Survival analysis of mice with double and single KO in cytokine receptor genes **(C)** or cytokine genes **(D)**, as indicated, after infection with VV_WR_. Mice that had lost more than 25% of the initial body weight were considered moribund and sacrificed.

**fig. S21.**
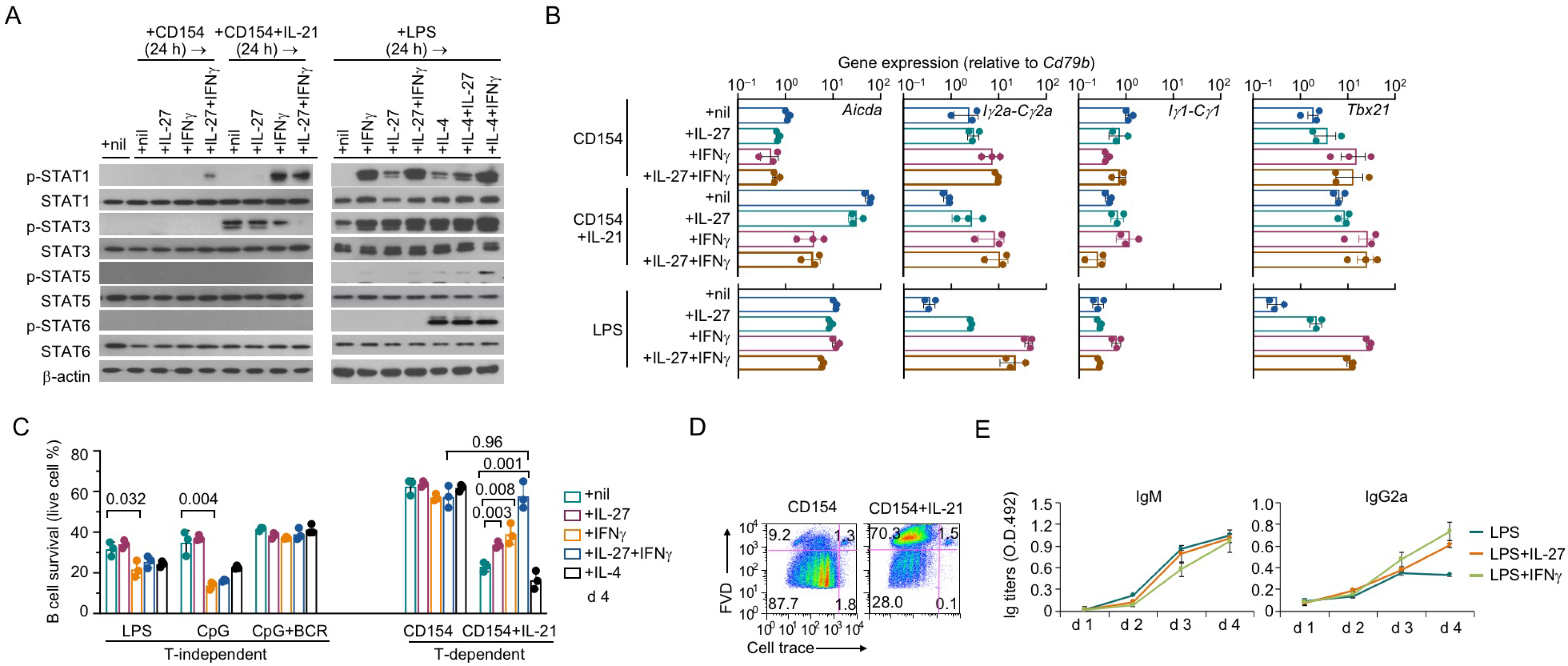
IL-27 and IFNγ regulate B cell differentiation. **(A)** Immunoblotting of phosphorylation of STAT1, STAT3, STAT5 and STAT6 in B cells stimulated with LPS (left), CD154 or CD154 plus IL-21 (right) for 24 h and then with different cytokines or cytokine combinations, as indicated, for 1 h. Representative of two independent experiments. **(B)** qRT-PCR analysis of gene induction in purified spleen B cells after stimulation for 24 h with CD154, CD154 plus IL-21 or LPS in the presence of different cytokines or cytokine combinations, as indicated. Data were normalized to *Cd79b* expression and expressed as the ratio to values of B cells stimulated with CD154 (top) or LPS (bottom). **(C)** Flow cytometry analysis of the proportion of live (7-AAD^-^) cells after B cells were stimulated with stimuli, as indicated, for 96 h. **(D)** Flow cytometry analysis of cell death (FVD^+^) after B cells were stimulated with CD154 plus IL-21, as indicated, for 96 h. **(E)** ELISA of IgM and IgG2a in B cells cultured for 96 h in the presence of LPS plus nil, IL-27 or IFNγ, as indicated.

**fig. S22.**
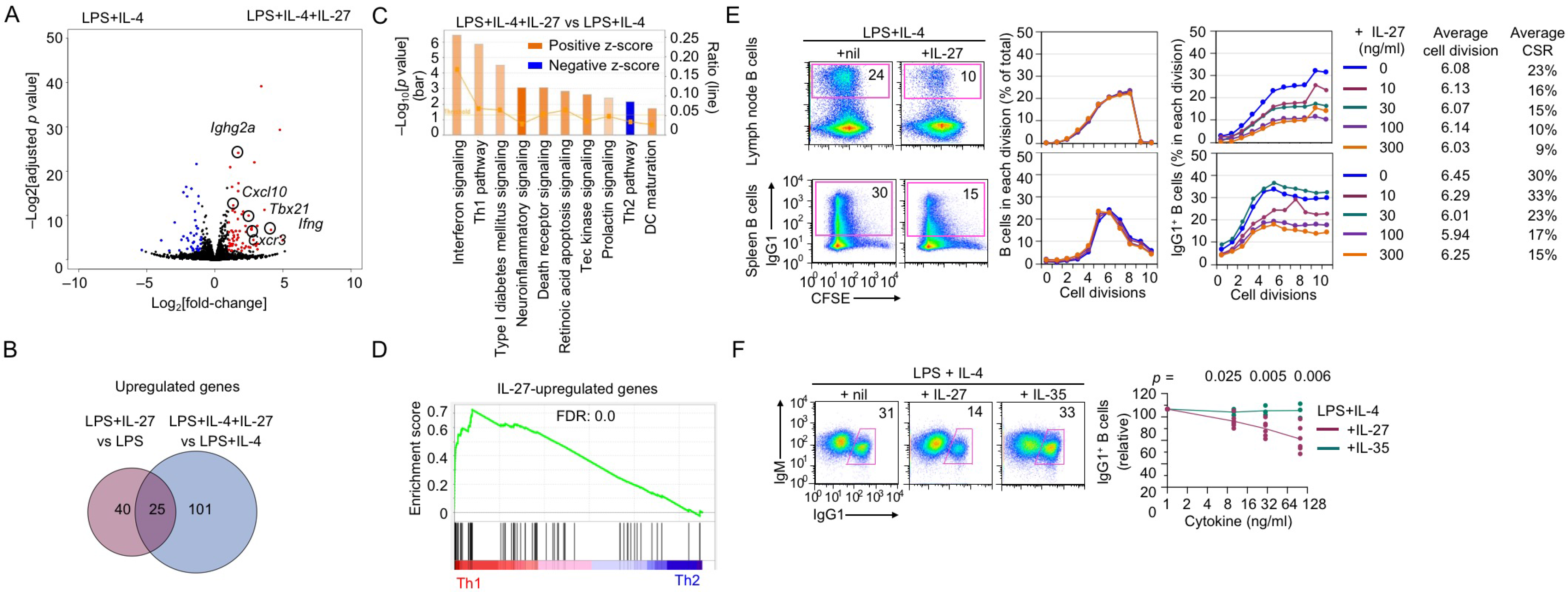
IL-27 promotes Th1 gene responses in B cells stimulated in the presence of IL-4 and inhibits CSR to IgG1. **(A)** Volcano plot depicting the induction of Th1 genes and IgG2a-related transcripts by IL-27 in B cells stimulated with LPS and IL-4 for 48 h. **(B)** Venn diagram depicting the genes upregulated by IL-27 (25) in both LPS-stimulated B cells and B cells stimulated with LPS and IL-4. **(C,D)** Ingenuity pathway analysis **(C)** and GSEA **(D)** of upregulation of the Th1 pathway genes and downregulation of the Th2 pathway genes by IL-27 in B cells stimulated with LPS and IL-4 for 48 h. (E) Flow cytometry analysis of proliferation and CSR to IgG1 in B cells isolated from the spleen or lymph nodes, labeled with CFSE and stimulated with LPS plus IL-4 in the presence of nil or different doses of IL-27 for 96 h. (F) Flow cytometry analysis of CSR to IgG1 in B stimulated with LPS plus IL-4 in the presence of nil or different doses of IL-27 or IL-35 for 96 h. Representative of two independent experiments.

**fig. S23.**
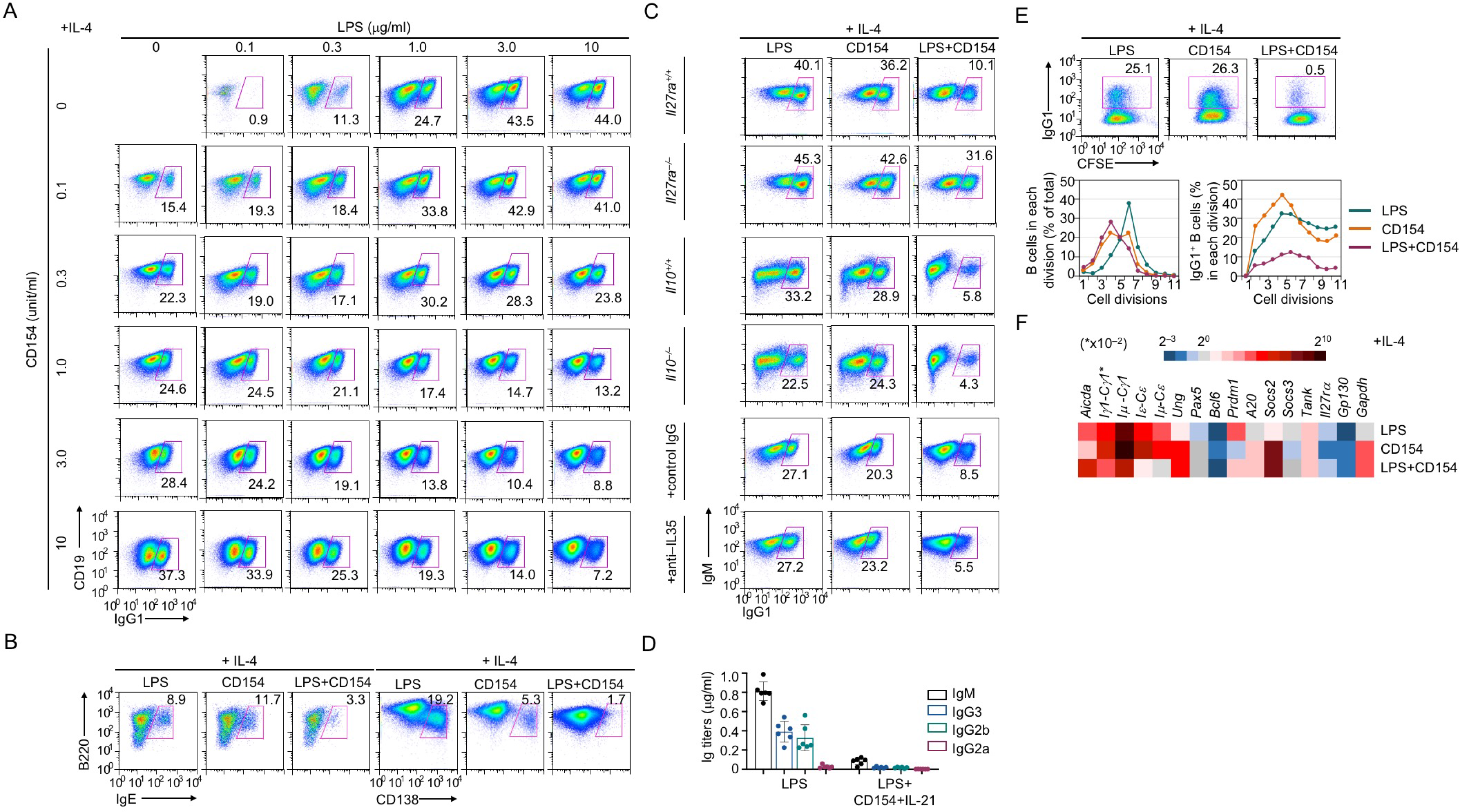
Inhibition of CSR and plasma cell differentiation in B cells induced to produce IL-27. **(A)** Flow cytometry analysis of CSR to IgG1 in purified B cells stimulated with LPS and CD154 at different doses, as indicated, plus IL-4 for 96 h. **(B)** Flow cytometry analysis of CSR to IgE and generation of plasmablasts/plasma cells in purified B cells stimulated with LPS and CD154 plus IL-4 for 96 h. Representative of tfour independent experiments. **(C)** Flow cytometry analysis of CSR to IgG1, as induced by LPS and CD154 plus IL-4 for 96 h, in B cells of different genotypes (top 4 rows) or treated with anti-IL-35 blocking Ab or a control IgG (bottom two rows). Representative of three independent experiments. **(D)** ELISA of Ig isotypes secreted by B cells stimulation with LPS (left) or LPS plus CD154 and IL-21 (to induce IL-27) for 96 h. **(E)** Flow cytometry analysis of proliferation and CSR to IgG1 in B cells labeled with CFSE and stimulated, as indicated, for 96 h (top), and depiction of the proportion of live B cells and IgG1^+^ B cells that had completed different numbers of divisions (bottom). **(F)** qRT-PCR analysis of gene expression in B cells stimulated for 48 h. Data were normalized to *Cd79b* expression and expressed as the ratio to values of freshly isolated B cells (0 h).

**fig. S24.**
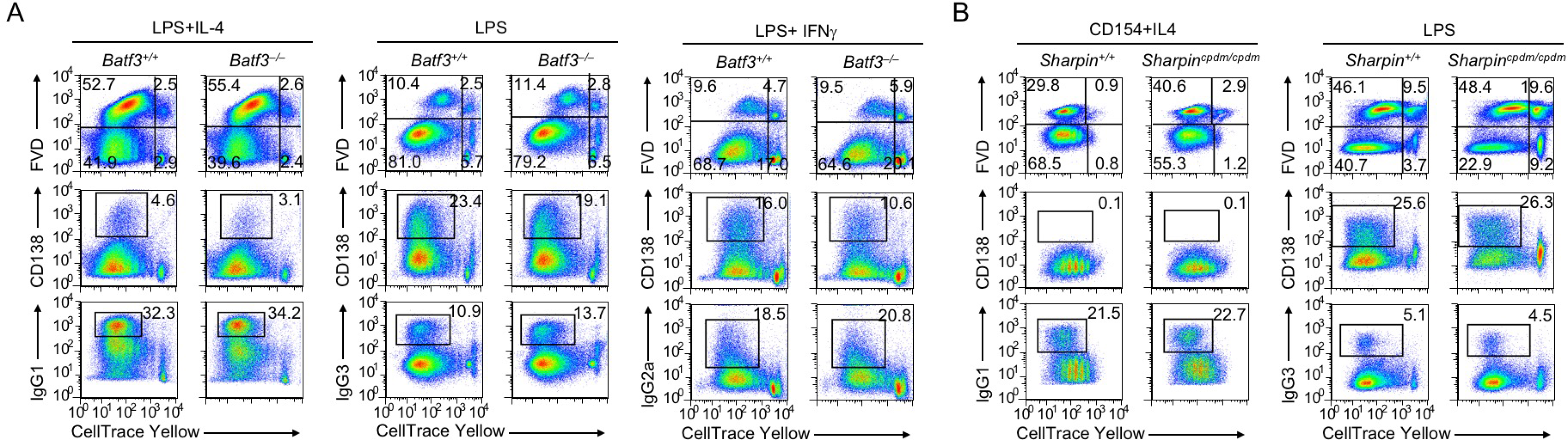
Normal proliferation, survival, CSR and plasma cell differentiation in B cells lacking *Batf3* or expressing *Sharpin^cpdm/cpdm^*. **(A)** Flow cytometry analysis of proliferation, survival, CSR to appropriate IgG isotypes and plasma cell differentiation in *Batf3^−/−^* and *Batf3^+/+^* B cells labeled with CellTrace Yellow and stimulated, as indicated, for 96 h. Representative of three independent experiments. **(A)** Flow cytometry analysis of proliferation, survival, CSR to appropriate IgG isotypes and plasma cell differentiation in *Sharpin^cpdm/cpdm^* and *Sharpin^+/+^* B cells labeled with CellTrace Yellow and stimulated, as indicated, for 96 h. Representative of four independent experiments.

**fig. S25.**
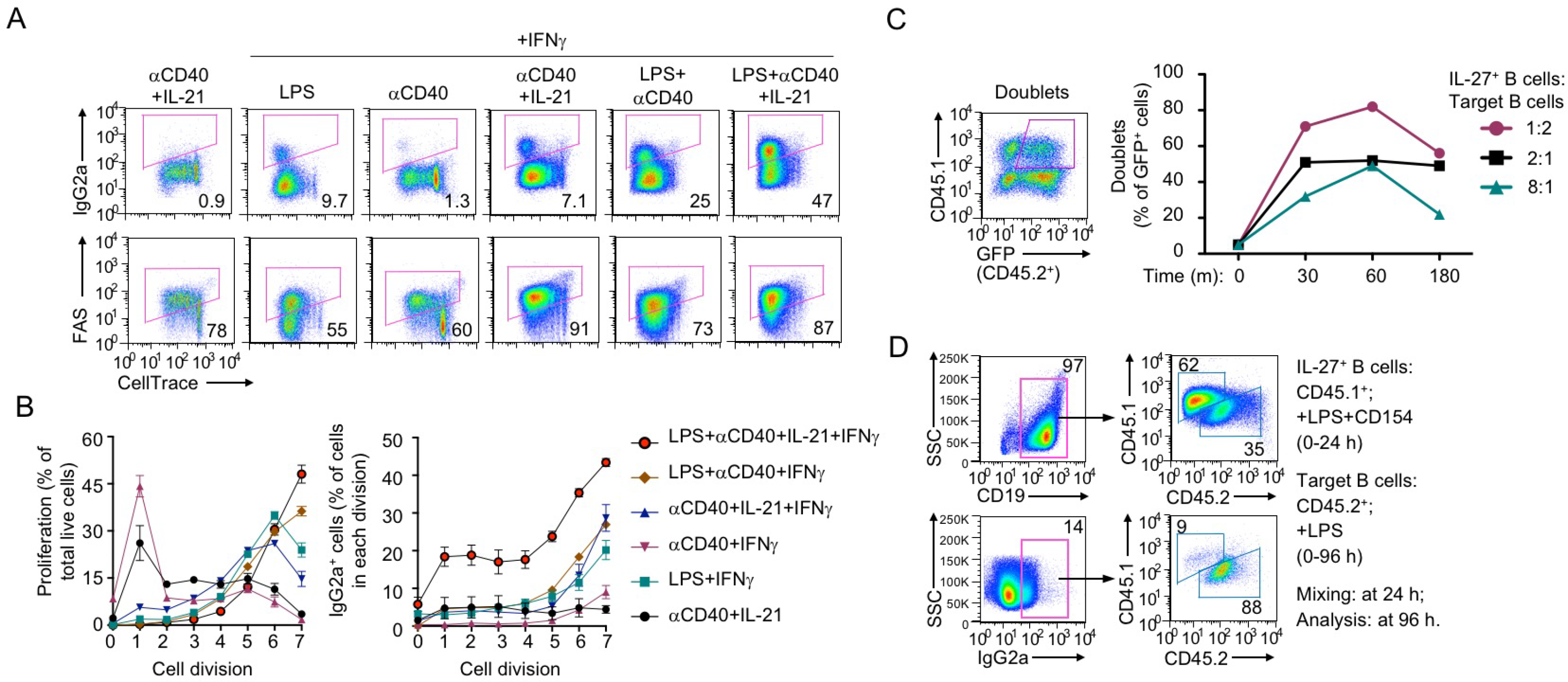
IL-27-producing B cells induce CSR to IgG2a in target B cells. **(A,B)** Flow cytometry analysis of CSR to IgG2a (**A,** top) and FAS expression **(A,** bottom), proliferation/survival (**B**, left) and CSR to IgG2a (**B**, right) in B cells labeled with CellTrace™ and stimulated, as indicated, for 96 h. Representative of two independent experiments. **(C)** Flow cytometry analysis of doublet formation by *Il27*-GFP^+^ B cells (indicating IL-27+ B cells), as generated by stimulation with LPS plus CD154 and IL-21 for 24 h, and their target B cells, i.e., B cells primed with LPS for 24 h, after being mixed at different ratios for 1 h. Representative of three independent experiments. (D) Flow cytometry analysis of CSR to IgG2a in IL-27-producing (CD45.1^+^) B cells, as generated *in vitro* by stimulation with LPS plus CD154 for 24 h, and (CD45.2^+^) B cells, as primed by LPS for 24 h, after being mixed and cultured for additional 72 h. Representative of three independent experiments.

**fig. S26.**
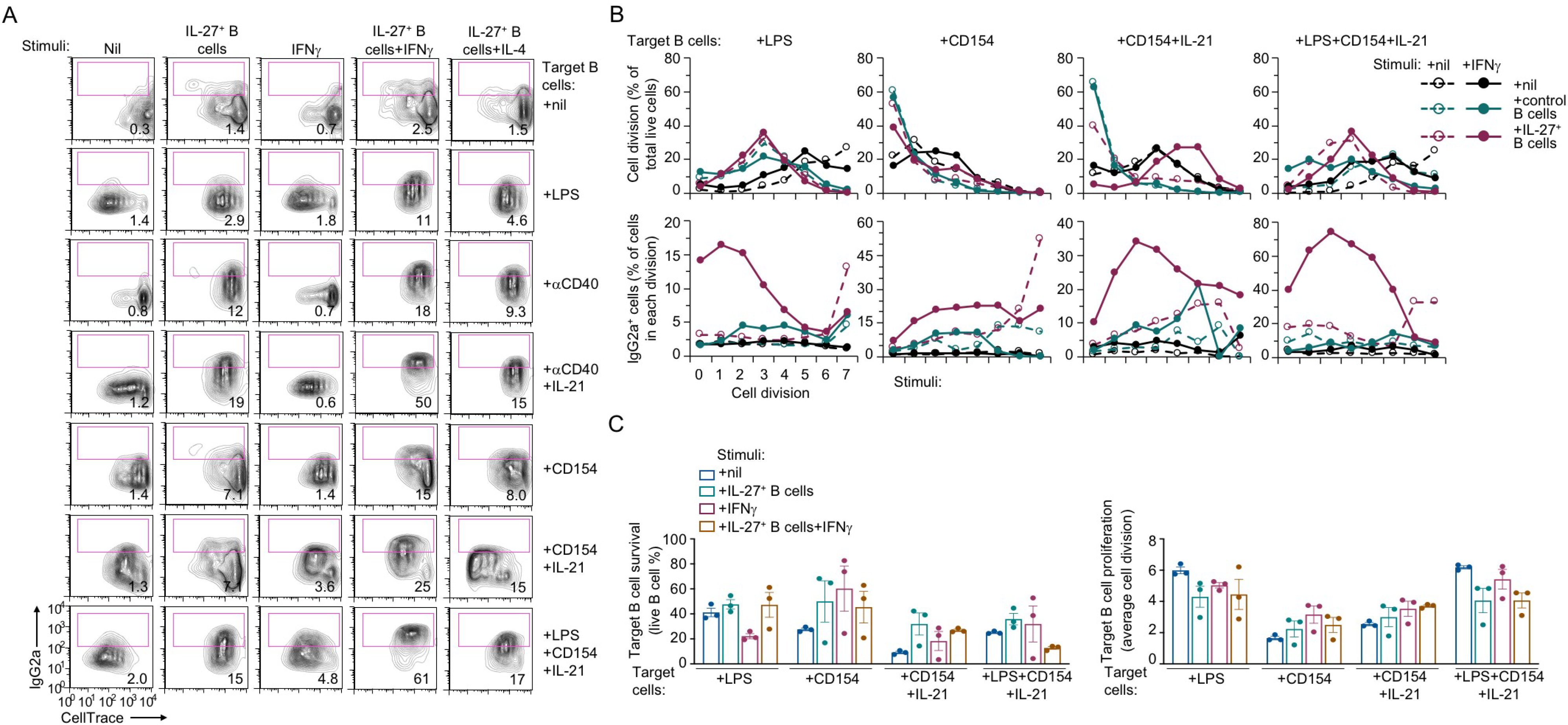
IL-27-producing B cells synergize with IFNγ to induce CSR to IgG2a in target B cells. **(A)** FACS analysis of proliferation and CSR to IgG2a in CD45.2^+^ target B cells labeled with CellTrace™ (to track proliferation) and activated, as indicated, 16 h before co-cultured for 72 h with nil, CD45.1^+^ IL-27^+^ B cells or “control” CD45.1^+^ B cells (generated by LPS stimulation for 36 h), in the presence of nil or IFNγ, as indicated. Representative of three independent experiments. **(B)** FACS analysis of proliferation and CSR to IgG2a in CD45.2^+^ target B cells, as in **(A)**. Depicted were the proportion of live B cells and lgG1^+^ B cells that had completed different numbers of divisions, as indicated. Representative of three independent experiments. **(C)** FACS analysis of survival and proliferation of CD45.2^+^ target B cells, as in **(A)** and **(B).** Depicted were the proportion of live B cells in target B cells (left; n=3) and the average numbers of divisions of target B cells (right; n=3).

