## Supplementary Tables for "BATF3-dependent induction of IL-27 in B cells bridges the innate and adaptive stages of the antibody response": Table S19 S20.docx

| **Table S19.** Antibodies. | | |
| --- | --- | --- |
| Flow Cytometry | | |
| α–mo/huEBI3-PE | eBioscience | Cat. # 12735842 (Clone ebic6) |
| α–moB220-Pacific Blue™ | Biolegend | Cat. # 103227 (RA3-6B2) |
| α–moCD11b-APC | BioLegend | Cat. # 101212 (Clone M1/70) |
| α–moCD11c-eFluor^®^ 450 | eBioscience | Cat. # 48-0114-82 (Clone N418) |
| α–moCD138-PE-Cy7 | BioLegend | Cat. # 142513 (Clone 281-2) |
| α–moCD19- PerCP-Cy5.5 | BioLegend | Cat. # 144605 (Clone 1D3) |
| α–moCD19-Pacific Blue™ | BioLegend | Cat. # 115523 (Clone 6D5) |
| α–moCD1d-BV510™ | BD Biosciences | Cat. # 563189 (Clone 1B1) |
| α–moCD3-Pacific Blue™ | BioLegend | Cat. # 100412 (17A2) |
| α–moCD38-PE-Cy7 | BioLegend | Cat. # 102718 (Clone T10) |
| α–moCD4-APC | BioLegend | Cat. # 100412 (GK1.5) |
| α–moCD40-PE | Biolegend | Cat. # 124610 (Clone 3/23) |
| α–moCD45-Pacific Blue™ | BioLegend | Cat. # 103126 (Clone 30-F11) |
| α–moCD45-PerCP | BioLegend | Cat. # 103130 (Clone 30-F11) |
| α–moCD8-PE | BioLegend | Cat. # 100412 (53-5.8) |
| α–moCD95-BV510™ | BD Biosciences | Cat. # 563646 (Clone Jo2) |
| α–moGL-7-PE | BioLegend | Cat. # 144607 (Clone GL7) |
| α–moGr-1-Pacific Blue™ | BioLegend | Cat. # 108430 (Clone RB6-8C5)  RB6-8C5) |
| α–moIgD-APC | BioLegend | Cat. # 405714 (Clone 11-26c.2a) |
| α–moIgG1-APC | BioLegend | Cat. # 406609 (Clone RMG1-1) |
| α–moIgG1-FITC | BioLegend | Cat. # 406606 (Clone RMG1-1) |
| α–moIgG2a-APC | BioLegend | Cat. # 407110 (Clone RMG2a-62) |
| α–moIgG2b-FITC | BioLegend | Cat. # 406705 (Clone RMG2b-1) |
| α–moIgG3-BV421™ | BD Biosciences | Cat. # 565808 (Clone R40-82) |
| α–moIgM-APC | BioLegend | Cat. # 406509 (Clone RMM-1) |
| α–moIgM-PE | BioLegend | Cat. # 406507 (Clone RMM-1) |
| α–moIL-27p28-APC | BioLegend | Cat. # 516906 (Clone MM27-7B1) |
| α–moIL-27p28-PE-Cy7 | BioLegend | Cat. # 516910 (Clone MM27-7B1) |
| α–moEBI3-PerCP | Novus Biologicals | Cat. # NBP203943 (Clone 5P10D3) |
| α–moIL-27Rα-PE | BD Biosciences | Cat. # 564337 (Clone 2918) |
| α–moκLight Chain-PE | Invitrogen | Cat. # MKAPPA04 |
| α–moCD45.1-BV510™ | BioLegend | Cat. # 110741 (Clone A20) |
| α–moCD45.2-PE-Cy7 | BioLegend | Cat. # 109830 (Clone 104) |
| α–moMHC Class II -PE | TONBO | Cat. # 50-5321 (Clone M5/114.15.2) |
| Immunofluorescence staining |  |  |
| α–moB220-FITC | BioLegend | Cat. # 103206 (Clone RA3-6B2) |
| α–moGL7 Alexa Flour^®^ 647 | BioLegend | Cat. # 144605 (Clone GL7) |
| α–moIL-27p28-APC | BioLegend | Cat. # 516906 (Clone MM27-7B1) |
| ELISA/ELISPOT | | |
| α–moIgM-UNLB | SouthernBiotech | Cat. # 1020-01 |
| α–moIgG-UNLB | SouthernBiotech | Cat. # 1030-01 |
| α–moIgA-UNLB | SouthernBiotech | Cat. # 1040-01 |
| α–moIgM-biotin | SouthernBiotech | Cat. # 1020-08 |
| α–moIgG1-biotin | SouthernBiotech | Cat. # 1070-08 |
| α–moIgG2a-biotin | SouthernBiotech | Cat. # 1080-08 |
| α–moIgG2b-biotin | SouthernBiotech | Cat. # 1090-08 |
| α–moIgG3-biotin | SouthernBiotech | Cat. # 1100-08 |
| α–moIgA-biotin | SouthernBiotech | Cat. # 1040-08 |
| α–moIgG-biotin | SouthernBiotech Technology | Cat. # 1030-08 |
| Other flow cytometry reagents | | |
| Fixable Viability Dye eFlour™ 780 | eBioscience | Cat. # 65-0865-14 |
| Fixable Viability Dye eFlour™ 506 | eBioscience | Cat. # 65-0866-14 |
| NP_23_-PE | Biosearch Technologies | Cat. # N-5070-1 |
| Immunoblotting/ChIP |  |  |
| α–moBATF3 | Abcam | Cat. # ab229727 |
| α–moβ-Actin | Cell signaling | Cat. # 3700 |
| α–mop-NF-κB p65 (Ser536) | Cell signaling | Cat. # 3033 |
| α–moNF-κB p65(D14E12) | Cell signaling | Cat. # 8242 |
| α–mop-STAT1 (Tyr701) (D4A7) | Cell signaling | Cat. # 7649 |
| α–moSTAT1 (D1K9Y) | Cell signaling | Cat. # 14994 |
| α–mop-STAT3 (Tyr705) (D3A7) | Cell signaling Technology | Cat. # 9145 |
| α–moSTAT3 (D1B2J) | Cell signaling | Cat. # 30835 |
| α–mop-STAT5 (Tyr694) (D47E7) | Cell signaling | Cat. # 4322 |
| α–moSTAT5 (D2O6Y) | Cell signaling | Cat. # 94205 |
| α–mop-STAT6 (Tyr641) | Cell signaling | Cat. # 9361 |
| α–moSTAT6 (D3H4) | Cell signaling | Cat. # 5397 |
| Goat α–rabbit IgG, HRP-linked | Cell signaling | Cat. # 7074 |
| α–moNF-κB p65 | Santa Cruz | Cat. # sc372 |

| **Table S20.** PCR Primers. | | | |
| --- | --- | --- | --- |
|  | | Forward Primer | Reverse Primer |
| Mouse genes | | | |
| *Aicda* | 5'-AGAAAGTCACGCTGGAGACC-3' | | 5'-CTCCTCTTCACCACGTAGCA-3' |
| *Batf* | 5'-CTGGCAAACAGGACTCATCTG-3' | | 5'-GGGTGTCGGCTTTCTGTGTC-3' |
| *Batf3* | 5'-CAGAGCCCCAAGGACGATG -3' | | 5'-GCACAAAGTTCATAGGACACAGC-3' |
| *Cd79b* | 5'-CCACACTGGTGCTGTCTTCC-3' | | 5'-GGGCTTCCTTGGAAATTCAG-3' |
| *Ebi3* | 5'- CGGTGCCCTACATGCTAAAT-3' | | 5'- GCGGAGTCGGTACTTGAGAG-3' |
| *Gapdh* | 5'-TTCACCACCATGGAGAAGGC-3' | | 5'-GGCATGGACTGTGGTCATGA-3' |
| *Iμ-Cγ1* | 5'-ACCTGGGAATGTATGGTTGTGGCTT-3' | | 5'-ATGGAGTTAGTTTGGGCAGCA-3' |
| *Iμ-Cγ2a* | 5'-ACCTGGGAATGTATGGTTGTGGCTT-3' | | 5'-GCTGGGCCAGGTGCTCGAGGTT-3' |
| *Iγ1-Cγ1* | 5'-TCGAGAAGCCTGAGGAATGTG-3' | | 5'-ATGGAGTTAGTTTGGGCAGCA-3' |
| *Iγ2a-Cγ2a* | 5'-GCTGATGTACCTACCGAGAGA-3' | | 5'-GCTGGGCCAGGTGCTCGAGGTT-3' |
| *Il10* | 5'- AGCCGGGAAGACAATAACTG-3' | | 5'-CATTTCCGATAAGGCTTGG-3' |
| *Il12a* | 5'- CATCGATGAGCTGATGCAGT-3' | | 5'- CAGATAGCCCATCACCCTGT-3' |
| *Il6* | 5'- GCTACCAAACTGGATATAATCAGGA-3' | | 5'- CCAGGTAGCTATGGTACTCCAGAA-3' |
| *P19* | 5'- CCAGCAGCTCTCTCTCGGAATC-3' | | 5'- TCATATGTCCCGCTGGTGC-3' |
| *P28* | 5'- CTCTGCTTCCTCGCTACCAC-3' | | 5'- GGGGCAGCTTCTTTTCTTCT-3' |
| *P40* | 5'- TTATGTTGTAGAGGTGGACTGG-3' | | 5'- TTTCTTTGCACCAGCCATGAGC-3' |
| *Prdm1* | 5'-GCTGCTGGGCTGCCTTTGGA-3' | | 5'-GGAGAGGAGGCCGTTCCCCA-3' |
| *Tbx21* | 5'- AACCGCTTATATGTCCACCCA-3' | | 5'- CTTGTTGTTGGTGAGCTTTAGC-3' |
